# Apple ripening is controlled by a NAC transcription factor

**DOI:** 10.1101/708040

**Authors:** Zoë Migicovsky, Trevor H. Yeats, Sophie Watts, Jun Song, Charles F. Forney, Karen Burgher-MacLellan, Daryl J. Somers, Yihi Gong, Zhaoqi Zhang, Julia Vrebalov, James G. Giovannoni, Jocelyn K. C. Rose, Sean Myles

**Affiliations:** Department of Plant, Food and Environmental Sciences, Dalhousie University, Faculty of Agriculture, Agricultural Campus, Truro, NS, Canada; Plant Biology Section, School of Integrative Plant Science, Cornell University, Ithaca, NY 14853 USA; Agriculture and Agri-Food Canada, Kentville, Nova Scotia, B4N 1J5, Canada; Vineland Research and Innovation Centre, Vineland Station, ON, L0R 2E0, Canada; College of Horticulture, South China Agriculture University, Guangzhou, China; Boyce Thompson Institute, Cornell University, Ithaca, NY 14853 USA; United States Department of Agriculture, Robert W. Holley Center, Cornell University, Ithaca, NY 14853 USA

**Keywords:** Apple, fruit ripening, NAC-domain transcription factor

## Abstract

Softening is a hallmark of ripening in fleshy fruits, and has both desirable and undesirable implications for texture and postharvest stability. Accordingly, the timing and extent of ripening and associated textural changes are key targets for improving fruit quality through breeding. Previously, we identified a large effect locus associated with harvest date and firmness in apple (*Malus domestica*) using genome-wide association studies (GWAS). Here, we present additional evidence that polymorphisms in or around a transcription factor gene, *NAC18.1*, may cause variation in these traits. First, we confirmed our previous findings with new phenotype and genotype data from ∼800 apple accessions. In this population, we compared a genetic marker within *NAC18.1* to markers targeting three other firmness-related genes currently used by breeders (*ACS1*, *ACO1*, and *PG1*), and found that the *NAC18.1* marker was the strongest predictor of both firmness at harvest and firmness after three months of cold storage. By sequencing *NAC18.1* across 18 accessions, we revealed two predominant haplotypes containing the single nucleotide polymorphism (SNP) previously identified using GWAS, as well as dozens of additional SNPs and indels in both the coding and promoter sequences. *NAC18.1* encodes a protein with high similarity to the NON-RIPENING (NOR) transcription factor, a regulator of ripening in tomato (*Solanum lycopersicum*). To test whether these genes are functionally orthologous, we introduced both *NAC18.1* transgene haplotypes into the tomato *nor* mutant and showed that both haplotypes complement the *nor* ripening deficiency. Taken together, these results indicate that polymorphisms in *NAC18.1* may underlie substantial variation in apple firmness through modulation of a conserved ripening program.

## Introduction

Despite their diverse structure, ontogeny, and biochemical composition, fleshy fruits from a taxonomically broad range of species undergo coordinated ripening processes that have many features in common. Ripening involves numerous physiological and biochemical changes that render the fruit attractive and nutritious for consumption by seed-dispersing animals, or human consumers in the case of crops. These changes include the accumulation of sugars, pigments, and flavor or aroma compounds, as well as a loss of flesh firmness due in large part to the controlled modification and depolymerization of cell wall polysaccharides (Wang et al., 2018). Processes involved in ripening are regulated by conserved and convergently evolved networks of transcription factors and hormones, such as ethylene in climacteric fruit where a respiratory burst occurs at the beginning of ripening (Lü et al., 2018).

Various aspects of fruit ripening have been particularly well-studied in tomato (*Solanum lycopersicum* L.) and the characterization of tomato ripening mutants has revealed a regulatory network consisting of transcription factors, hormones, and epigenetic modifications (Giovannoni et al., 2017). Among the best studied ripening-related transcription factors in tomato is NON-RIPENING (NOR), a NAC [No apical meristem (NAM), *Arabidopsis* transcription activation factor (ATAF), Cup-shaped cotyledon (CUC)] domain transcription factor (Giovannoni et al., 2004) expressed early in ripening (Shinozaki et al., 2018). NAC genes comprise one of the largest plant-specific families of transcription factors, with specific members regulating development, defense, and senescence (Mathew and Agarwal, 2018). While all NAC genes share a conserved DNA-binding (NAC) domain, specific functional clades are defined in terms of their more variable domains, particularly the C-terminal transcriptional regulatory region. These domains can act directly as transcriptional activators, or can facilitate interaction with other transcription factors in order to fine-tune transcriptional control. NAC transcription factors have been implicated in ripening phenotypes in diverse species including tomato (Kumar et al., 2018), melon (Ríos et al., 2017), banana (Shan et al., 2012), peach (Pirona et al., 2013) and apricot (García-Gómez et al., 2019).

Apple (*Malus* x *domestica* Borkh.) fruit exhibit extensive variation in the extent and timing of ripening and softening. In previous genome-wide association studies (GWAS) of 689 apple accessions, we found a single nucleotide polymorphism (SNP) on chromosome 3 within the coding sequence of a NAC transcription factor, *NAC18.1* (MD03G1222600 in the GDDH13 v1.1 reference genome (Daccord et al., 2017)), associated with harvest date and firmness (Migicovsky et al., 2016). This SNP results in an aspartate (D) to tyrosine (Y) mutation at a highly conserved position of the *NAC18.1* amino acid sequence and we refer to this putatively causal SNP as D5Y. Subsequently, GWAS of three additional germplasm collections confirmed the association between the genomic region containing *NAC18.1* and ripening time (Urrestarazu et al., 2017; Larsen et al., 2019; Jung et al., 2020). Thus, the *NAC18.1* gene is a strong candidate for mediating apple ripening time and firmness, and the D5Y SNP may be of utility for marker-assisted breeding.

Due to a prolonged juvenile phase, it is particularly challenging for apple breeders to evaluate fruit quality traits. As a result, in recent years considerable effort has been invested in developing molecular markers that can be used to select for fruit quality traits at the seedling stage. In particular, markers in three genes are currently used by apple breeders to select for desirable fruit texture. The first is *1-AMINOCYCLOPROPANE-1-CARBOXYLATE SYNTHASE 1* (*ACS1*), which encodes the ripening-associated isoform of an ethylene biosynthesis gene. The *ACS1-2* allele contains a retrotransposon insertion thought to confer low ethylene production and longer shelf life in ‘Fuji’ and other apple cultivars homozygous for this allele (Sunako et al., 1999; Harada et al., 2000). The second gene corresponds to another enzyme involved in ethylene biosynthesis, *AMINOCYCLOPROPANE-1-CARBOXYLATE OXIDASE 1* (*ACO1*), which has a similar reduced-functionality allele (Costa et al., 2005). Finally, *POLYGALACTURONASE 1* (*PG1*) encodes an enzyme that hydrolyzes pectin polysaccharides in the cell wall and middle lamella and thus has been implicated in affecting apple fruit firmness (Atkinson et al., 2012). Several apple *PG* alleles have been identified (Costa et al. 2010). Apple firmness was identified as one of the five most important traits for genomics-assisted breeding (Laurens et al., 2018) and markers for the desirable alleles of *ACS1*, *ACO1*, and *PG1* are widely available (Baumgartner et al., 2016).

The ripening of climacteric fruits, like apples, is regulated by the plant hormone ethylene (Grierson, 2013). All three firmness-related genes described above are ethylene-dependent: their expression is mediated by ethylene after the initiation of the climacteric process and is repressed by exposure to the ethylene inhibitor 1-methylcyclopropene (1-MCP) (Brummell and Harpster, 2001; Dandekari et al., 2004; Costa et al., 2005, 2010; Tadiello et al., 2016; Di Guardo et al., 2017). While *PG1* has been associated with apple firmness using GWAS (Kumar et al., 2013; Di Guardo et al., 2017), the discovery of *ACS1*, *ACO1* and *PG1* markers was driven largely by linkage mapping studies of bi-parental families (Costa et al., 2010; Longhi et al., 2012, 2013; Bink et al., 2014; Sadok et al., 2015). These studies did not consistently detect all three loci, which may have been due to limited sample sizes, lack of segregation between parents, and/or differences in phenotyping methods. Subsequent testing has revealed either low, or no power for these markers to predict firmness phenotypes, raising doubt about their utility in marker-assisted breeding (Costa et al., 2013; Nybom et al., 2013; McClure et al., 2018; Chagné et al., 2019).

Here we extend our previous work and report an evaluation of the role of *NAC18.1* in apple ripening and softening. First, we tested the utility of the three established firmness-related markers (*ACO1*, *ACS1* and *PG1*) and the *NAC18.1* D5Y marker to predict harvest date, firmness and softening during storage across more than 800 diverse apple accessions. Second, we determined the effect of ethylene and 1-MCP on the expression levels of all four genes. Third, we sequenced the *NAC18.1* gene from a subset of apple cultivars to discover potentially causal alleles in linkage disequilibrium (LD) with D5Y. Finally, we heterologously expressed the *NAC18.1* gene in the tomato *non-ripening (nor)* mutant to test whether it functions as a component of a conserved fruit ripening program via complementation.

## Materials and Methods

### Germplasm sources

Apple samples taken for this study included three different sources of germplasm. The majority of the samples were from Canada’s Apple Biodiversity Collection (ABC), an orchard located in Kentville, Nova Scotia, which contains 1,113 apple accessions. A comprehensive description of the ABC is provided in Watts et al. (submitted). Briefly, the ABC is a diverse germplasm collection planted in duplicate in an incomplete block design, which includes 1 of 3 standards per grid, allowing for correction of positional effects using a REstricted Maximum Likelihood (REML) model, described in Migicovsky et al. (2017). Samples from the ABC were used for phenotyping of harvest date and firmness, as well as genotyping of texture-related genetic markers.

Additional apple samples were taken from the United States Department of Agriculture (USDA) apple germplasm collection and the Cornell apple breeding program in Geneva, New York. Samples from Geneva were used for sequencing of *NAC18.1*.

Lastly, ‘Golden Delicious’ apples from a commercial orchard in Nova Scotia were harvested before the climacteric stage from 2004-2007, as described previously (Yang et al., 2013, 2016). Samples from this previous work were used to test the expression levels of genes of interest.

### Apple phenotyping

In 2017, we evaluated harvest date for 1,348 trees and fruit firmness for 1,328 trees within the ABC. Due to the diversity of apples within the collection, a variety of methods were used to determine the appropriate time to harvest. First, we observed if the tree had dropped fruit or, for red apples, if the fruit were a deep red color. Next, a sample apple was taken from each tree and touched to assess firmness, tasted to assess starch and sweetness, cut in half to check browning of seeds, and then sprayed with iodine solution to assess starch content (Blanpied and Silsby, 1992a). Fruit were deemed mature and ready to harvest at a starch-iodine index of 6 (Blanpied and Silsby, 1992b).

Once harvested, the fruit were evaluated for firmness. We recorded the firmness (kg/cm^2^) of 5 fruit per tree using a penetrometer with a 1 cm diameter (Fruit Texture Analyzer, GS-14, Güss Manufacturing). A small section of skin was removed using a vegetable peeler, and each fruit was placed on the penetrometer platform so that the piston entered the middle of the apple where the skin had been removed. The data were automatically recorded into a spreadsheet. After these measurements were taken, 5 additional fruit from each accession (combined across trees/replicates) were placed in cold storage (5°C). Fruit was removed from storage after 3 months and was left at room temperature for 24 hours before being evaluated again for firmness using the same method. Firmness after storage and the percent change in firmness from harvest to post-storage were both calculated.

Harvesting fruit from the ABC orchard often lasted more than one day and so differences in harvest date within a week reflect the time required to harvest the orchard, rather than meaningful biological differences. As a result, we recorded harvest dates as the Monday of each week for all trees harvested throughout the week. We used the “lmer” function in the R package lme4 (Bates et al., 2015) to fit a REML model for harvest date and firmness at harvest. Next, we calculated the least squares mean using the “lsmeans” function in the lsmeans R package (Lenth, 2016), resulting in one value per accession. After running the REML model, we had 862 unique accessions with harvest dates and 859 accessions with firmness at harvest measurements. Due to the number of fruits available and storage capacity, replicates were combined from multiple trees prior to storage and so it was not possible to fit a REML model for firmness after storage or change in firmness measurements, which both had sample sizes of 535 unique accessions. We calculated the correlation between each of the phenotypes using Pearson correlation tests and the “ggpairs” function in the GGally R package (Schloerke et al., 2020).

### Texture-related genetic markers

DNA was extracted from leaf tissue collected from the ABC orchard using silica columns, quantified using PicoGreen (Thermo) and normalized to a concentration of 20 ng µL^-1^. Genotyping was conducted using PCR and high resolution melting (HRM) on a LightScanner HR384 (BioFire). Primers are listed in Supplemental Table S1.

Genotyping of the *PG1* SNP marker was based on the GenBank sequence L27743.1, where the T allele is favorable at position 437 and the G allele is unfavorable and leads to increased softening during storage (Costa et al., 2010). Three allelic combinations of the observed indel, *ACS1-1/1*, *1-1/2* and *1-2/2*, have been associated with high, medium, and low ethylene production, respectively (Sunako et al., 1999; Harada et al., 2000; Oraguzie et al., 2004; Costa et al., 2005). The exact position of the indel from *ACS1-2* (GenBank: AB010102.1) is 1,320 to 1,483 bp (163 bp), and from *ACS1-1* (GenBank: AY062129.1) is 4,500 to 4,525 bp (25 bp). The size difference between alleles is 138 bp. The *ACO1* marker involves an unfavorable 62 bp insertion in the third intron of *ACO1* (Costa et al., 2005). The third intron spans from 1,083 bp to 1,300 bp of the GenBank sequence Y14005.1 and the indel is found from position 1,297 bp to 1,358 bp. The D5Y mutation in *NAC18.1* is a nonsynonymous SNP at position 30,698,039 on chromosome 3, according to reference genome version GDDH13 v1.1 <u>(Daccord et al. 2017)</u> and is associated with both harvest date and firmness (Migicovsky et al., 2016). The desirable C allele encodes an aspartic acid (D) at the fifth amino acid position of the NAC18.1 protein, while the undesirable A allele encodes a tyrosine (Y). The names and IDs of samples, their phenotypes and their genotypes are provided in Table S2.

### Phenotypic variance explained by texture-related markers

To determine the proportion of phenotypic variance explained by each marker of interest (ACO1, ACS1, PG1, and NAC18.1) we used a type 2 ANOVA from the ‘car’ package in R (Fox and Weisberg, 2018) with the markers encoded as co-dominant. To determine the phenotypic variance explained by each marker after accounting for harvest date, we also performed a type 2 ANOVA including the four markers and harvest date as factors. The results of the models were visualized using the “geom_tile” function in the ggplot2 R package (Wickham, 2016). We determined the association between each marker and each phenotype using Spearman’s rank correlation test. We visualized the results using the “geom_boxplot” function in ggplot2 in R (Wickham, 2016).

### Sequencing of NAC18.1

DNA was isolated from leaves sampled from 18 apple accessions growing in Geneva, NY. A 2.3 kb amplicon including the *NAC18.1* gene and ∼800 bp of upstream sequence was amplified by PCR using primers NAC18F2 and NAC18R2 (Table S1) and Phusion® High-Fidelity PCR Master Mix with HF Buffer (NEB). PCR product size and purity was confirmed by agarose gel electrophoresis, and the remaining product was purified using a DNA Clean & Concentrator kit (Zymo Research). The resulting DNA fragment was cloned into the plasmid pMiniT 2.0 and transformed into *E. coli* using the NEB® PCR Cloning Kit (NEB).

Individual colonies were selected for complete sequencing of the cloned amplicon using the primers NAC18F2, NAC18F3, NAC18F4, NAC18R1, and NAC18R2 (Table S1). For accessions homozygous for the D5Y SNP, the NAC18.1 amplicon from a single clone was sequenced. For heterozygous accessions, two clones representing each D5Y allele were selected based on partial sequencing of the D5Y region, followed by complete sequencing of the 2.3 kb amplicon, as described above. The nucleotide sequences were aligned using MUSCLE (Edgar, 2004) and used to construct a maximum-likelihood phylogenetic tree in MEGA7 (Kumar et al., 2016).

### Gene expression analyses of apple fruit

Gene expression levels of *ACO1*, *ACS1*, *PG1*, and *NAC18.1* were evaluated using q-PCR with and without treatment of ethylene and 1-methylcyclopropene (1-MCP), using methods described previously in (Yang et al., 2013, 2016). Briefly, in 2004 and 2005, ‘Golden Delicious’ fruit were harvested from a commercial orchard before the climacteric stage. Ethylene gas (36 µL L^-1^) was applied for 24 hours at 20°C to initiate ripening while control fruits were stored for 24 hours at 20°C without ethylene. All fruit were then stored at 20°C for 21 days, with sampling occurring at day 0, 7, 13, and 21 of each year. In 2006 and 2007, ‘Golden Delicious’ were once again harvested at the pre-climacteric stage. Apples were either treated with 1-MCP (1 µL L^-1^ of EthylBloc, 0.14%, Rohm and Haas Company) for 12 hours at 20°C in a sealed container, or stored at 20°C for 12 hours without 1-MCP. Fruits were then stored at 20°C, with sampling occurring on day 0, 7, 14, and 22.

Total RNA was extracted from frozen apple tissues using a hot borate method with some modification in the extraction buffer, as described in (Yang et al., 2016). RNA extracts were treated with DNase I using a DNA-free Kit following the manufacturer’s recommendations (Applied Biosystems). First-strand cDNA synthesis was performed using 2 µg DNase I-treated total RNA. The oligonucleotide primers used for real-time quantitative qRT-PCR analysis were designed from sequence information in NCBI (Table S1). Conditions for all PCR reactions were optimized as previously described (Yang et al., 2013) and efficiency values for each gene are shown in Table S3. Two reference genes, *MdActin* and *MdUBI*, were used in the real-time qPCR analysis to normalize the expression patterns. Samples from day 0 (assigned an arbitrary quantity of “1”) were used as a calibrator to calculate relative quantities (Yang et al., 2016). The experimental design was a balanced randomized block design with random effects of year, plate, and row (on plate). The fixed effects were treatments for both ethylene (for example, control (Day 7) and ethylene (Day 7)) and 1-MCP experiments (such as control (Day 7) and 1-MCP (Day 7)).

### Molecular cloning and plant transformation

The hypothetical coding sequences (CDSs) corresponding to the consensus sequence of the “A” and “C” haplotypes of *NAC18.1* were synthesized as a double-stranded DNA (gBlock) by Integrated DNA Technologies (IDT), with 20 bp of flanking sequences added to both ends to facilitate Gibson Assembly of the *NAC18.1* sequences between the *Asc*I and *Pac*I restriction sites of pMDC32 (Curtis and Grossniklaus, 2003). Constructs were assembled using the NEBuilder HiFi DNA Assembly Cloning Kit (NEB) and their integrity verified by Sanger sequencing. Plasmids were transformed into *Agrobacterium tumefaciens*, which was used to transform tomato (*S. lycopersicum*) cotyledon explants (Van Eck et al., 2019) derived from the tomato *nor* mutant in the cv. Ailsa Craig background (LA3770, Tomato Genetics Resource Center, https://tgrc.ucdavis.edu/). A total of 10 T0 plants were recovered per construct and plants derived from two independent transformation events per construct were selected for further characterization in the T1 generation.

### Transgenic tomato characterization

Fruit were harvested when visually ripe, or in the case of the *nor* mutant, at the equivalent age as ripe cv. Ailsa Craig fruit (the near isogenic wild-type control), as determined by tagging of flowers at anthesis. The color of the fruit surface was measured using a CR-400 Chroma Meter (Konica Minolta), and fruit were weighed, photographed, and dissected. Pericarp tissue was frozen in liquid nitrogen and stored at -80°C. Frozen tissue was ground to a fine powder and RNA extracted using a modified version of the protocol described in Chang et al. (1993). Briefly, approximately 400 mg of tissue was added to a preheated (80°C) two-phase system consisting of 500 µL of water-saturated phenol and 500 µL of extraction buffer (100 mM Tris [pH 8.0], 25 mM EDTA, 2 M NaCl, 2% CTAB, 2% PVP, and 2% [v/v] beta-mercaptoethanol). The mixture was vortexed and incubated for 5 min at 65°C and then cooled to room temperature before extracting and precipitating RNA, as previously described (Chang et al., 1993).

RNA was treated with RNase-free DNase (Promega) and used for cDNA synthesis with RNA to cDNA EcoDry™ Premix with Oligo dT primer (Takara Bio). The cDNA was used as a template for quantitative PCR using Luna Universal qPCR Master Mix (NEB) and a Viia7 real-time PCR instrument (Life Technologies/ABI). Gene-specific primers are listed in Table S1. Quantification used the ΔC_t_ method with *RIBOSOMAL PROTEIN L2* (*RPL2*) as a reference gene, and statistical significance of the ΔC_t_ values was tested by a one-way ANOVA followed by Tukey’s HSD test.

## Results

### Evaluation of texture-related markers

Fruit were harvested over a 65 day period and their firmness at harvest (N = 859) and after 3 months of cold storage (N = 535) was found to differ by 7-fold across the apple accessions from the ABC. We observed a strong relationship between harvest date and firmness: late-harvested apples were firmer both at harvest (R^2^ = 0.25, p < 1 x 10^-15^) and after storage (R^2^ = 0.24, p < 1 x 10^-15^), and they also softened less during storage (R^2^ = 0.086, p = 4.53 x 10^-12^) (Figure S1). Firmness at harvest was also significantly correlated with firmness after storage (R^2^ = 0.54, p < 1 x 10^-15^; Figure S1). Post-harvest storage resulted in significant softening: on average, apples lost 40 percent of their firmness during 3 months of cold storage (Table S2).

We tested the utility of four genetic markers to predict firmness-related phenotypes and found that the *NAC18.1* marker outperformed the other three for both firmness at harvest and firmness after storage. However, softening (the loss in firmness during storage) was best predicted by the *PG1* marker. Our results suggest that the markers in *ACO1* and *ACS1* have little to no predictive power for firmness-related traits across diverse apple germplasm (Figure 1; Figure S2). When we performed the same analysis with harvest date as a factor, harvest date accounted for 10.9% and 14.6% of the variation in firmness at harvest and firmness after storage, respectively. In this model, the amount of phenotypic variance in firmness at harvest explained by *NAC18.1* was reduced from 18.19%, when harvest date was not included in the model, to 2.64% (Figure 1; Figure S3).

**Figure 1:**
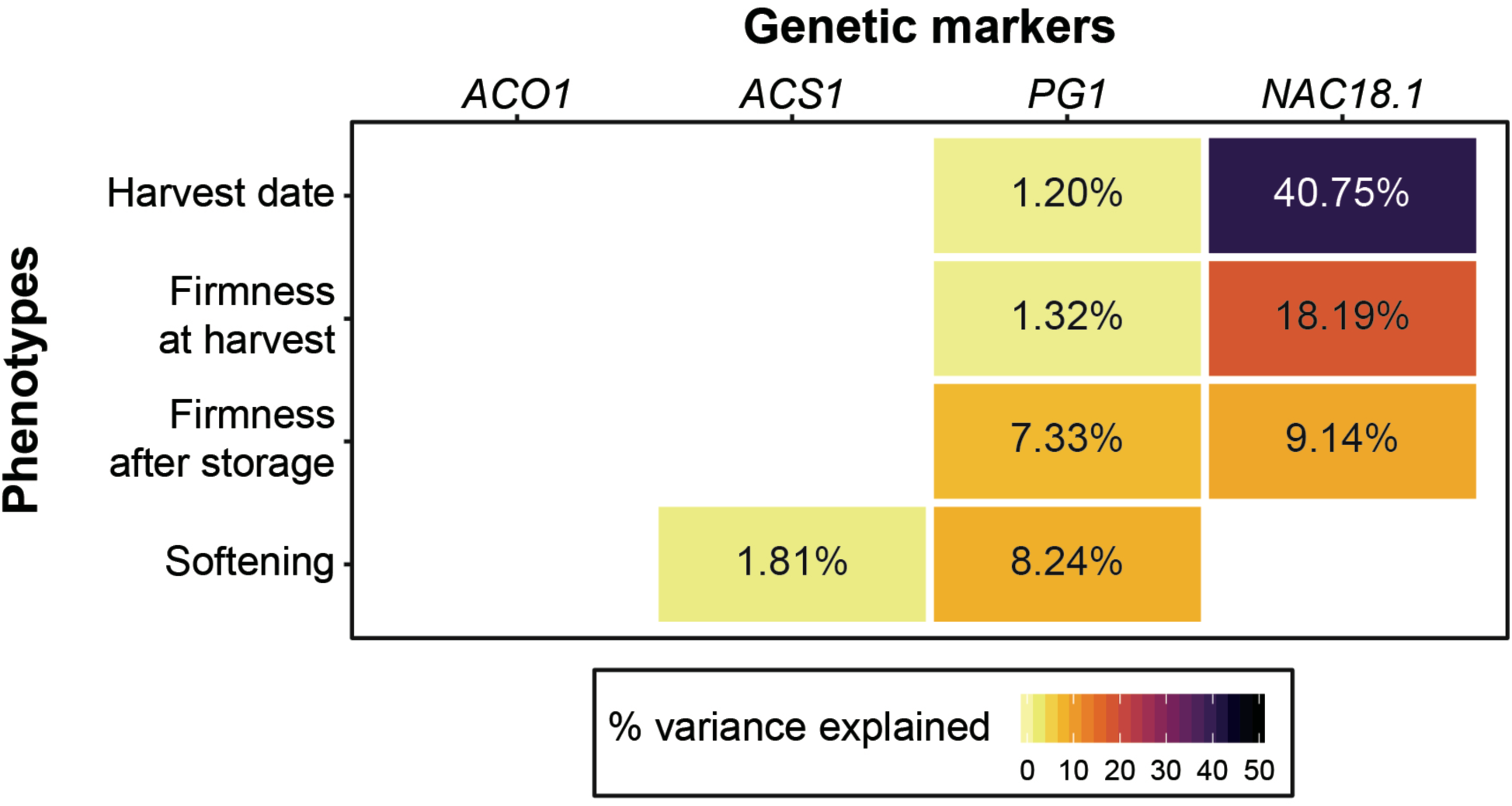
The utility of four genetic markers for predicting harvest date and firmness-related apple phenotypes. For each trait, the percent variance explained for each of the four markers is shown in cases where a significant effect was detected from a type 2 ANOVA. A type 2 ANOVA was also performed with the four genetic markers and harvest date as a co-factor (Figure S3).

The genotypes of the four texture-related genetic markers across the nine most popular apple cultivars sold in the USA in 2018 (Home - USApple 2021) are presented in Figure 2. All nine cultivars were homozygous for the desirable (firm) C allele of *NAC18.1*, while only 2 to 4 of the cultivars had homozygous desirable genotypes for the other markers. Among the top cultivars, only ‘Fuji’, released in 1962, was homozygous for desirable alleles at all markers, while ‘McIntosh’, initially discovered in 1811 and commercially released in 1870, was homozygous for undesirable alleles at all markers except for *NAC18.1*. ‘Empire’, an offspring of ‘McIntosh’ and ‘Red Delicious’ released in 1966, was homozygous for the desirable *NAC18.1* alleles like its parents, but inherited undesirable alleles from ‘McIntosh’ and thus was heterozygous for *ACS1* and *ACO1*, and homozygous for undesirable *PG1* alleles.

**Figure 2:**
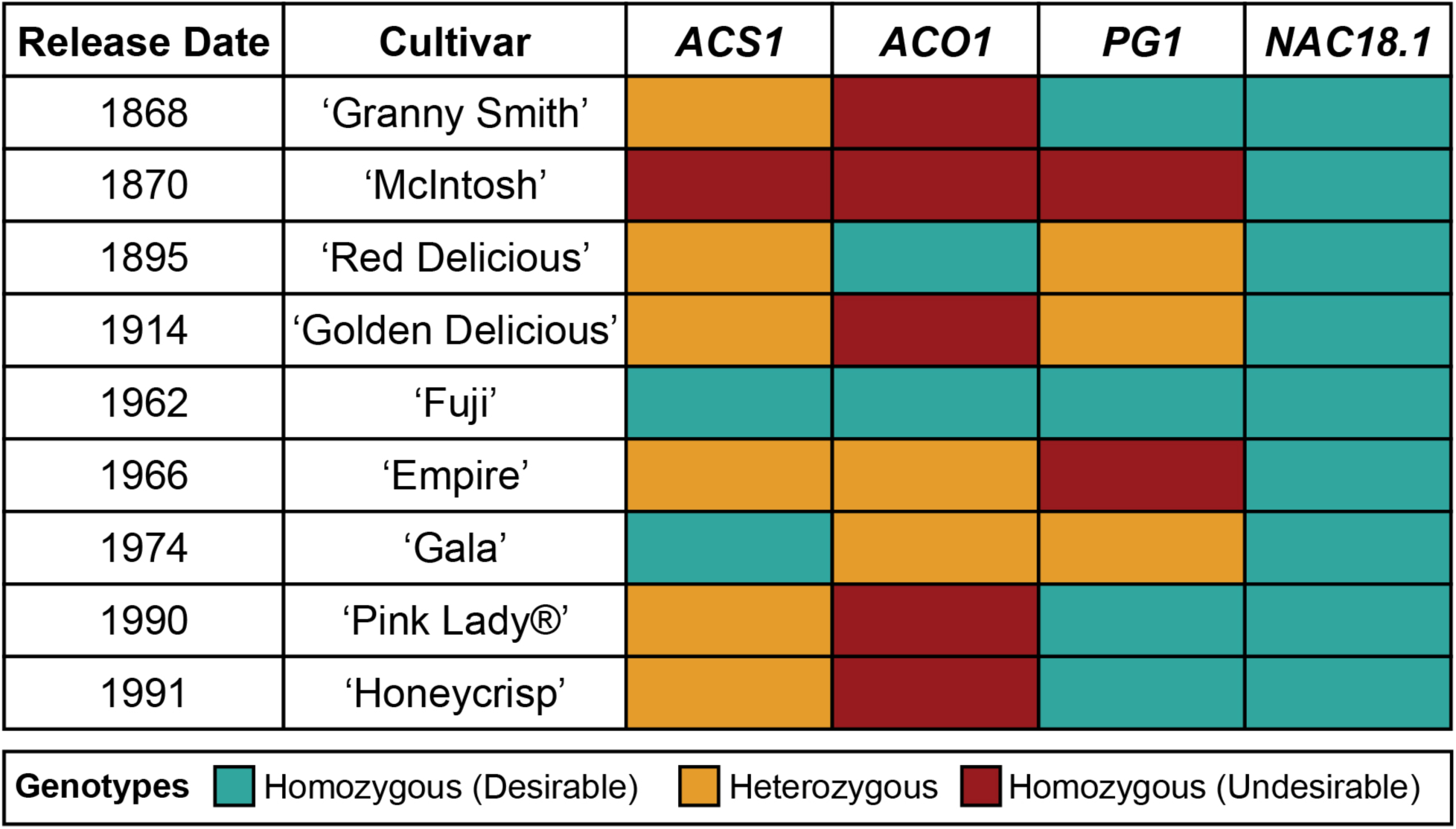
Genotypes of four texture-related genetic markers across the top 9 apple cultivars sold in the United States (U.S. Apple Association). The “desirable” allele for each marker is defined as the allele that has been reported to lead to firmer apple texture.

Using q-PCR, we evaluated the expression of the four candidate firmness genes across a three week period and observed that transcript levels of *ACO1*, *ACS1* and *PG1* genes were up-regulated in response to treatment with ethylene and suppressed following exposure to the ethylene-inhibitor, 1-MCP. *NAC18.1* transcript levels, however, remained unaffected by exposure to ethylene and 1-MCP (Figures S4 and S5).

### Resequencing of NAC18.1

DNA sequences from 24 *NAC18.1* haplotypes confirmed the expected D5Y genotype in all individuals and revealed a number of additional SNPs and indels within both coding and non-coding regions of *NAC18.1* (Table S4). A multiple sequence alignment and subsequent phylogenetic analysis indicated two major clades, corresponding to the D5Y A and C alleles (Figure 3A). In addition to the amino acid change resulting from D5Y, several additional amino acid changing polymorphisms were observed between sequences from D5Y genotypes and the reference sequence of *NAC18.1*. For example, near the site of the D5Y polymorphism, all “A” haplotypes also had a 12 nucleotide insertion that introduced the amino acid sequence QPQP (Figure 3B, Figure S6).

**Figure 3:**
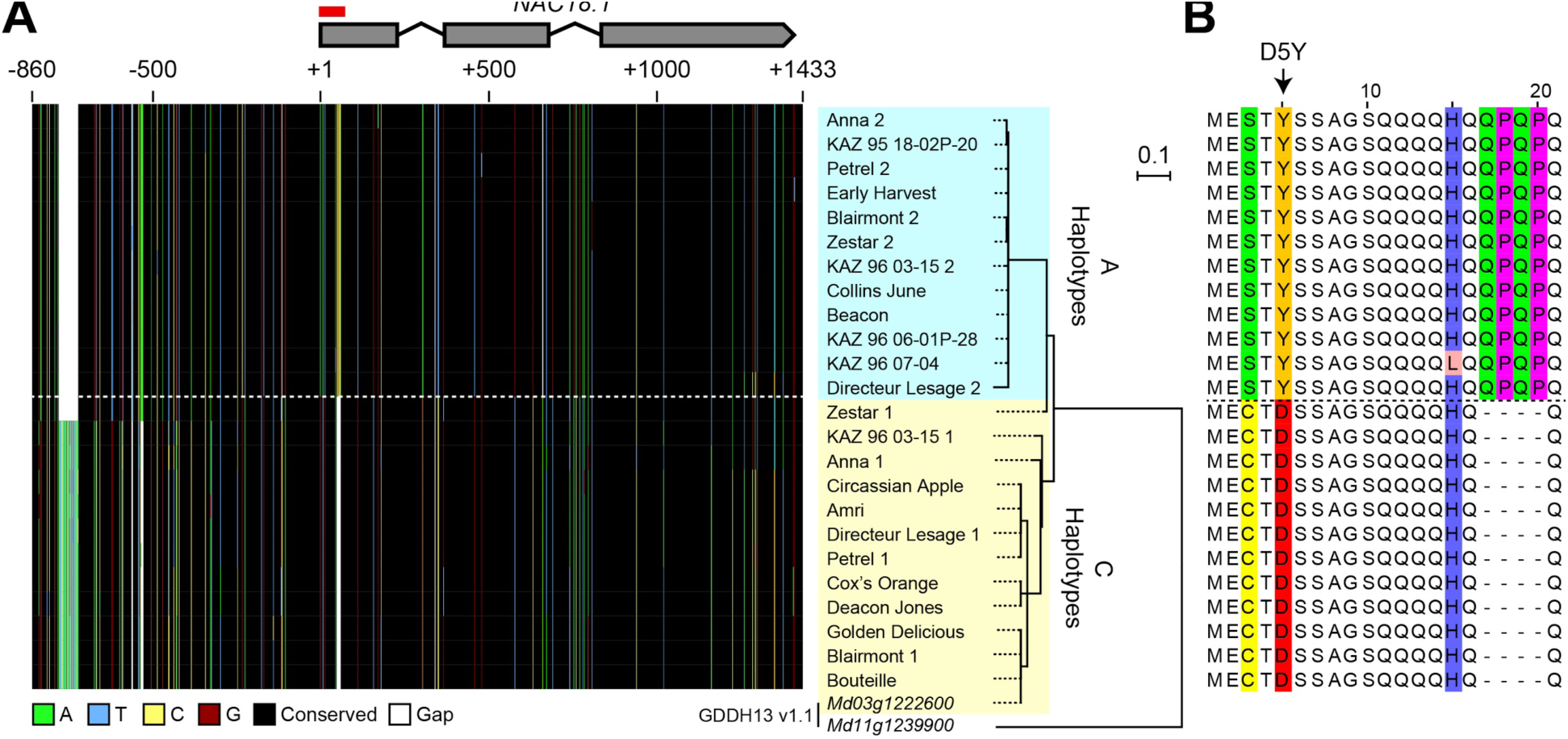
Polymorphisms within the *NAC18.1* gene. (A) Maximum likelihood phylogenetic tree of 24 *NAC18.1* sequences from 18 apple cultivars and the reference genome (GDDH13 v1.1) sequences of *NAC18.1* (Md03g1222600) and its closest homolog (Md11g1239900). A graphical representation of the 2.3 kb multiple sequence alignment is also shown. (B) Amino acid sequence alignment of the N-terminal region of *NAC18.1,* illustrating additional variation in the coding sequence in strong LD with the D5Y variant. The complete amino acid sequence alignment is provided in Figure S6.

### Transgenic complementation of the tomato non-ripening (nor) mutant

A BLAST search of predicted proteins from the ITAG 3.20 tomato genome (https://solgenomics.net/) using *NAC18.1* as a query identified only two matching sequences: *Solyc10g006880* (*NOR*) and *Solyc07g063420* (*NOR-LIKE 1*). To test the functional conservation of *NAC18.1* and *NOR*, we introduced constructs individually conferring constitutive expression of each of the *NAC18.1* haplotype CDSs into the tomato *nor* mutant. Two independent lines for each construct were characterized in the T1 generation with respect to their ability to rescue the ripening deficiency of the *nor* mutant. In contrast to *nor*, fruit from all four lines changed color at maturity, although internal fruit color change did not occur to the same extent as observed in a WT control (Figure 4A). To complement this qualitative phenotypic assessment, we also conducted quantitative colorimetry of the surface of the fruit (Figure 4B). Fruit from all the transgenic lines exhibited a significant increase in the *a\** (green-red) component of color space relative to the *nor* mutant, although only *NAC18.1^A^ #6* achieved similar *a\** levels to WT.

**Figure 4:**
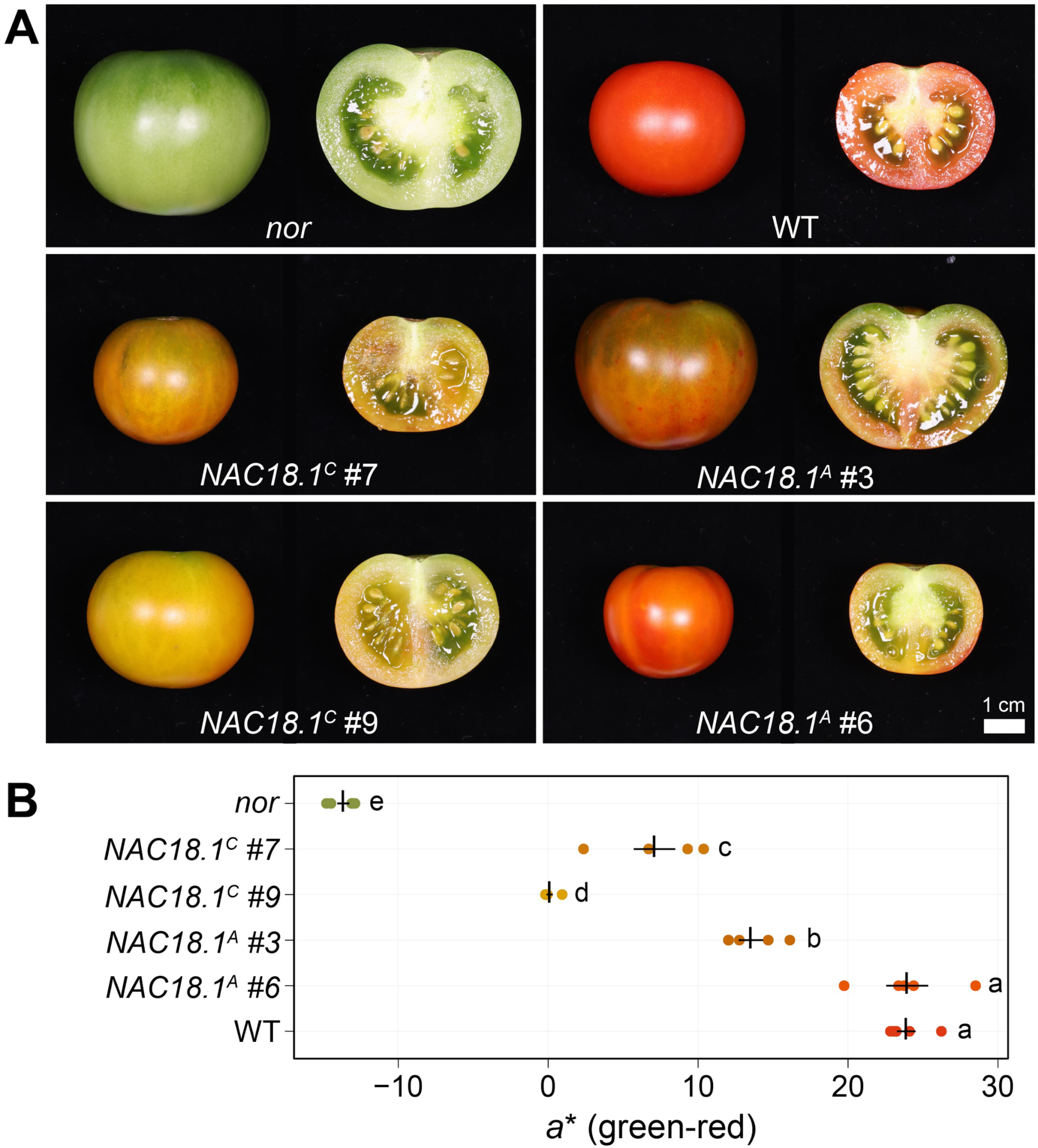
Transgenic complementation of the tomato *nor* mutant using *NAC18.1* transgene restores ripening. (A) Mature fruit of the tomato *nor* mutant and four independent T1 transgenic lines constitutively expressing either of two alleles of the *NAC18.1* transgene, *NAC18.1^C^* and *NAC18.1^A^* and isogenic WT control (cv. Ailsa Craig). (B) Quantitative colorimetry of the fruit surface of *nor,* the transgenic fruit and WT fruit. The *a*\* component (green-red axis) is shown, with the mean ± SE superimposed in black over the raw values (*n* = 5) in a color approximating the external color of the fruit. Genotypes not sharing a letter (a-e) are statistically distinct by one-way ANOVA and Tukey’s HSD test (*p* < 0.05).

The different degrees to which ripe fruit color was restored in the transgenic plants might be a consequence of either the different alleles of *NAC18.1*, or of different levels of transgene expression in each line. To address this, we analyzed the expression level of *NAC18.1* using qRT-PCR primers designed to target both *NAC18.1* alleles. Expression levels of the *NAC18.1* transgene were not statistically different between each independent line (*p* > 0.5) (Figure S7A). Next, we measured the expression level of several genes associated with tomato ripening physiology: *PHYTOENE SYNTHASE 1 (PSY1)* encodes an enzyme in an early stage of the carotenoid synthesis pathway, which is responsible for the production of red pigments during fruit ripening (Fray and Grierson, 1993), and its expression is impaired in the *nor* mutant (Osorio *et al*., 2011); *POLYGALACTURONASE 2* (*PG2*) encodes an enzyme catalyzing pectin depolymerization associated with fruit softening (Biggs and Handa, 1989); and *1-AMINOCYCLOPROPANE-1-CARBOXYLATE SYNTHASE 2* (*ACS2*) encodes an enzyme that synthesizes 1-aminocyclopropane-1-carboxylate, the immediate precursor of ethylene (Nakatsuka et al., 1998).

Expression of all three marker genes was enhanced in the *NAC18.1* transgenic lines relative to the *nor* mutant control, although not to the same extent as observed in WT ripe fruit (Figure S7B-D). Visual color change was consistent with the expression of the carotenoid biosynthetic gene *PSY1* (Figure S7B). Gene expression analysis further confirmed the induction of genes involved in ripening-associated cell wall remodeling (Figure S7C) and ethylene synthesis (Figure S7D). In contrast to the consistent level of *NAC18.1* expression observed in each line, the marker genes were more variable in their expression levels between lines. A similar pattern was observed for all marker genes, with the *NAC18.1^C^* #9 line showing the smallest induction of marker gene expression relative to *nor*. In the case of *PG2* and *ACS2*, the difference in expression in *NAC18.1^C^* #9 was not statistically significant relative to *nor* (*p* = 0.09 and 0.18, respectively). Consistent with these results, fruit from this line also exhibited the lowest amount of red color development (Figure S7B). Taken together, these results indicate that a canonical ripening program can be induced in the tomato *nor* mutant through the heterologous expression of either apple *NAC18.1* allele.

## Discussion

Genomics-assisted breeding has tremendous potential in perennial crops, such as apple, where a lengthy juvenile phase and large plant size make phenotyping at the adult stage time-consuming and expensive. However, the genetic markers used for culling progeny at the seedling stage must accurately predict the trait of interest at the adult stage in order for genomics-assisted breeding to be effective and cost-efficient (Luby and Shaw, 2001). Apple texture has been repeatedly identified as a key breeding target because of consumer demand for crisp, firm apples that retain their desirable texture during storage (Harker et al., 2003; Yue et al., 2013; Laurens et al., 2018). Markers for three genes (*ACS1*, *ACO1*, and *PG1*) are widely used in genomics-assisted apple breeding programs to predict firmness. However, GWAS for firmness suggests a single major effect locus for firmness at the *NAC18.1* gene (Migicovsky et al., 2016; Urrestarazu et al., 2017; McClure et al., 2018; Larsen et al., 2019). Here we have presented further evidence that whereas *ACS1*, *ACO1* and, *PG1* are weak predictors of firmness at harvest in genetically diverse apples, *NAC18.1* is a strong predictor and may serve a functional role in the fruit ripening pathway.

Single-marker correlation tests between each of the four markers and four phenotypes revealed statistically significant (*P* < 0.05) associations in every case except one (Figure S2). This result was not surprising: many genome-wide markers are expected to be correlated with firmness-related phenotypes since the genetic structure of our apple population is strongly correlated with these traits. For example, without correcting for the effects of population structure, 39% and 17% of genome-wide SNPs were significantly associated (*P* < 0.05) with harvest date and firmness, respectively, in the USDA collection, which is genetically identical to most of the population studied here (Migicovsky et al., 2016). Single-marker association tests do not account for population structure using genome-wide markers, as is customarily done when performing GWAS. Thus, we are only able to compare the relative power of each marker to predict these phenotypes. We found that the D5Y marker in *NAC18.1* had a 3 to 14 times greater effect on firmness at harvest and harvest date than the markers in *ACO1*, *ACS1* and *PG1* (Figure S2). We assessed the combined effects of the markers on these phenotypes using type 2 ANOVA and demonstrated that the D5Y marker in the *NAC18.1* gene is a far stronger predictor of harvest date and firmness at harvest than the markers for *ACS1*, *ACO1* and *PG1.* For predicting firmness after storage, however, *NAC18.1* performs only slightly better than *PG1*, while softening during three months of cold storage is best predicted by the marker in *PG1* (Figure 1). It is worth noting that a recent GWAS found that variation in firmness loss may not be due to variation within *PG1* but rather at a neighboring ethylene response factor (ERF) gene, *ERF* (MDP0000855671) (McClure et al., 2018). *ERF* may have been missed by previous linkage mapping studies due to a lack of mapping resolution compared to GWAS. Future fine mapping is required to determine if markers at *ERF* serve as superior predictors of softening during storage compared to the widely used *PG1* marker.

In our study, by far the strongest association observed for any marker-phenotype combination was between harvest date and the D5Y marker in *NAC18.1*: 41% of the variance in harvest date is accounted for by *NAC18.1* (Figure 1). On average, accessions homozygous for the late-ripening *NAC18.1* allele are harvested 29 days later than accessions homozygous for the early-ripening allele, which represents nearly half of the 65-day harvest season (Figure S2). This observation is consistent with strong GWAS signals for harvest date and ripening period discovered in and around the *NAC18.1* gene (Migicovsky et al., 2016; Urrestarazu et al., 2017; Larsen et al., 2019; Jung et al., 2020). The D5Y marker was also the strongest predictor of firmness at harvest and firmness after storage (Figure 1). We observed strong positive correlations between harvest date and both firmness at harvest (R^2^ = 0.25, p < 1 x 10^-15^) and firmness after storage (R^2^ = 0.24, p < 1 x 10^-15^; Figure S1), consistent with previous work showing that early-ripening apples tend to be softer (Vincent, 1989; Johnston et al., 2002; Oraguzie et al., 2004, 2007; Nybom et al., 2013; Chagné et al., 2014). When harvest date is included as a factor in the type 2 ANOVA instead of a response variable, it is the strongest predictor of firmness both at harvest and after storage (Figure S3). Thus, an apple’s firmness is largely determined by its harvest date. Therefore, markers that predict harvest date, such as the D5Y marker in *NAC18.1*, will be more effective for breeding than the other firmness-related markers tested here. This conclusion is in agreement with previous work suggesting that screening for *ACO1*, *ACS1* and *PG1* is not cost effective and that a marker for harvest date is of greater value to improving firmness via marker-assisted breeding (Nybom et al., 2013).

The best predictor of softening during storage was the marker in *PG1*, which accounted for 8% of the variance. The *PG1* marker was also significantly associated with firmness after storage, accounting for 7% of the variance in addition to the 9% accounted for by *NAC18.1* (Figure 1). Most apples are consumed after being stored. Thus, the most relevant phenotype for apple quality from the perspectives of both consumers and breeders is firmness after storage. The firmness of an apple after storage is a consequence of how firm it was when harvested and how much firmness loss it experienced during storage. Firmness loss, or softening, in climacteric fruit like apple is regulated by ethylene, and previous studies have shown that *ACO1*, *ACS1* and *PG1* transcripts increase when exposed to ethylene (Atkinson et al., 1998; Wakasa et al., 2006; Mann et al., 2008; Zhu and Barritt, 2008; Costa et al., 2010; Longhi et al., 2012; Di Guardo et al., 2017). We confirmed this result by showing that expression of these three genes is up-regulated by ethylene and down-regulated by the ethylene inhibitor, 1-MCP (Figures S4 and S5). The expression of *NAC18.1*, however, was unaffected by treatments with ethylene and 1-MCP. NAC transcription factors in apple are primarily involved in the regulation of growth and development (Li et al., 2020) and previous work has suggested that ethylene may not be required for on-tree apple ripening (Lau et al., 1986; Blankenship and Unrath, 1988). Our results suggest that genetic variation at the *NAC18.1* locus affects fruit firmness before harvest via a ripening pathway that is independent of ethylene, while softening during storage is ethylene-dependent and influenced by variation in or near *PG1*.

In the context of a marker-assisted breeding program, our results suggest that apple texture could be improved by selecting for the D5Y marker in *NAC18.1*. However, the potential for this marker to improve apple texture should be considered in light of several factors. First, we measured firmness with a penetrometer, but we recognize that our measures are only a proxy for the texture desired by consumers, which may be better captured using consumer panels and/or other mechanical devices that better mimic the chewing process (Costa et al., 2011). Second, all 9 of the most popular cultivars in the USA tested here are homozygous for the *NAC18.1* allele associated with late-harvested, firm apples (Figure 2). Out of 1,056 accessions genotyped, 696 (66%) are homozygous for the desirable *NAC18.1* allele, and its high frequency suggests this allele may already be under selection by breeders. Indeed, a recent population genetic analysis found evidence of positive selection for the desirable *NAC18.1* allele (Migicovsky et al., 2021). As a result, this allele may not segregate in many breeding populations. This may be the reason that *NAC18.1* was only identified as a firmness locus via GWAS in diverse populations, and not in numerous bi-parental breeding populations, emphasizing the need for germplasm collections that maintain diverse populations (Migicovsky et al. 2019). Third, highly aromatic apple cultivars are generally less firm, and selection for firmness using the D5Y marker may therefore result in lower production of volatile organic compounds, and thus apples with less pronounced aromas (Song and Bangerth, 1996; Farneti et al., 2017). Finally, selection for firmness using the D5Y marker will also select for late-harvested cultivars, which could result in a future excess of commercial cultivars with a compressed harvest window near the end of the harvest season, risking fruit loss to late season weather and reducing the opportunity to pick and sell fruit throughout the season. A more thorough understanding of the relationship between harvest date and firmness may lead to novel ways to break the correlation between these two phenotypes and to thereby enable the development of new apple cultivars with desirable firmness attributes, whose harvest dates are spread throughout the harvest season.

While the D5Y mutation in *NAC18.1* is a strong functional candidate variant for apple firmness, we discovered numerous DNA sequence variants in *NAC18.1*, often in perfect LD with D5Y, that are also putatively functional (Figure 3). The magnitude of DNA polymorphism at this locus, and the multitude of putatively causal variants, is consistent with the results of Larsen et al. (2019), who found 18 SNPs and 2 indels within *NAC18.1* across 11 apple accessions. Our analysis of *NAC18.1* sequences revealed a number of mutations that are candidates for causal association with ripening phenotypes. For example, we observed an insertion of four amino acids (QPQP) 11 amino acids downstream of the D5Y mutation in all A haplotypes upstream of the NAC DNA binding domain. Glutamine-rich sequences are common in eukaryotic transcription factors, and polymorphisms in these motifs have been shown to alter the activity of transcriptional activators (Atanesyan et al., 2012). On the other hand, we also observed several SNPs resulting in amino acid changes within the DNA binding domain and C-terminal transcriptional activator domain of *NAC18.1*. Given the pattern of LD we observed across *NAC18.1* (Figure 3), there remains the possibility that the GWAS signals at *NAC18.1* are driven by a causal variant outside of the *NAC18.1* coding sequence that acts independently of the *NAC18.1* gene. The identification of a causal polymorphism within the coding sequence or promoter of *NAC18.1* will require additional high-resolution genetic mapping and/or transcriptome analysis efforts.

Since *NAC18.1* is a homolog of the well-known ripening gene *NOR* in tomato, we used tomato as a model to explore the role of *NAC18.1* in the ripening process. We observed enhanced ripening in the tomato *nor* mutant following heterologous expression of either of the *NAC18.1* alleles (Figure 4). Although the expression level of the *NAC18.1* transgene was comparable across transgenic lines (Figure S7A), the level of ripening marker induction in the distinct lines was significantly different (Figure S7B-D). We had hypothesized that an allele-specific trend in our transgenic experiments would indicate that coding sequence variants are responsible for the association of *NAC18.1* with variation in apple ripening. While our results cannot confirm an allele-specific trend, they do verify that both alleles of *NAC18.1* encode functional proteins that can promote ripening.

In tomato, *nor* was first identified as a spontaneous mutation in an heirloom cultivar (Tigchelaar et al., 1973). While this *nor* allele exhibits recessive behavior and has been assumed to confer a complete loss-of-function, due to a 2 nucleotide deletion resulting in a truncated protein (Giovannoni et al., 2004), it was recently demonstrated that it is actually a dominant negative allele (Gao et al., 2019; Wang et al., 2019). These studies used CRISPR/Cas9 to generate *bona fide* null alleles of *nor*, which showed evidence of ripening relative to the spontaneous *nor* allele, although to a lesser extent than WT fruit. In light of these studies, we speculate that the action of the *NAC18.1* transgenes was likely attenuated by the dominant-negative activity of the spontaneous *nor* allele used in our work. Further heterologous characterization of *NAC18.1* would likely benefit from the use of null *nor* mutants, or double mutants of *nor* and *nor-like 1* (Gao et al., 2018). This may allow for more precise quantitative comparisons between alleles in order to resolve whether differences in the coding sequence of *NAC18.1* confer different degrees of ripening.

Although we are unable to conclude whether polymorphisms in coding or regulatory sequences affect the activity of *NAC18.1* in apple, an increasing body of evidence generated using tomato as a model system indicates that coding sequence polymorphisms of *NOR* can influence firmness and timing of ripening. For example, the ‘Alcobaça’ tomato cultivar has firm fruit, delayed ripening, and long shelf life that is conferred by the *alcobaça* mutation, and a recent study has shown that *ALCOBACA* is allelic with *NOR*, and that the *alcobaça* allele of *NOR* contains a valine to aspartate mutation at position 106 within the NAC domain (Kumar et al., 2018). An additional complete loss of function allele of *NOR* was also found in another long shelf life tomato cultivar (Kumar et al., 2018).

In conclusion, the results presented here provide evidence that *NAC18.1* is involved in apple ripening via an ethylene-independent mechanism, and that genetic variation in and/or near the *NAC18.1* gene influences apple ripening. The present study lays the groundwork for future efforts to compare the effect of different *NAC18.1* haplotypes and determine the precise causal variant(s) underlying this agriculturally important gene. Ultimately, we envision that the identification of precise causal variants for ripening in apple could be exploited using gene editing for fast and efficient introgression of desirable texture phenotypes.

## Data availability

The authors declare that all data supporting the findings of this study are available within the paper and its supplementary information files.

## Acknowledgments

We thank Susan Brown (Cornell University), Benjamín Gutiérrez (USDA-ARS Plant Genetics Resource Unit), and C. Thomas Chao (USDA-ARS Plant Genetics Resource Unit) for apple leaf tissue samples from orchards of the Cornell apple breeding program and the USDA apple germplasm collection. Transgenic tomato plants were generated by the Boyce Thompson Institute Biotechnology Center (https://btiscience.org/). ZM was supported by National Science Foundation Plant Genome Research Program 1546869. TY and JR were supported by a grant from the USDA-ARS Multistate (NE-1336) project, NYC-184821. This research was supported in part by funding the National Sciences and Engineering Research Council of Canada (SM) and A-Base funding (NOI-1238) from Agriculture and Agri-Food Canada (JS). The authors gratefully acknowledge the farm services team at the Kentville Research and Development Centre for maintaining the apple collection studied here.

## Supplemental Figures

**Figure S1:**
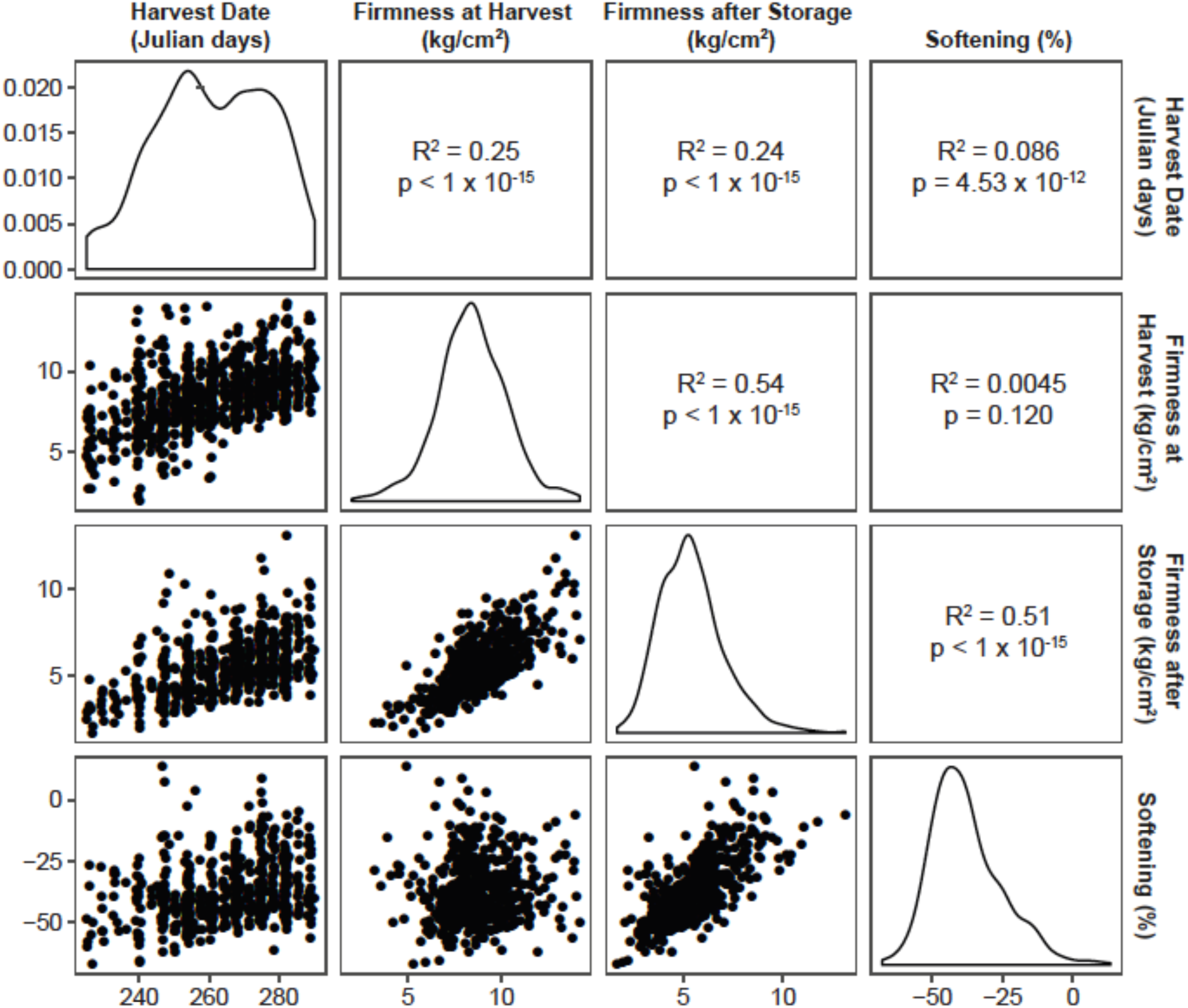
Correlations among phenotypes. The distributions of each phenotype are shown as well as dot plots of comparisons between each pair of phenotypes. The results of a Pearson correlation test are provided for each pairwise comparison.

**Figure S2:**
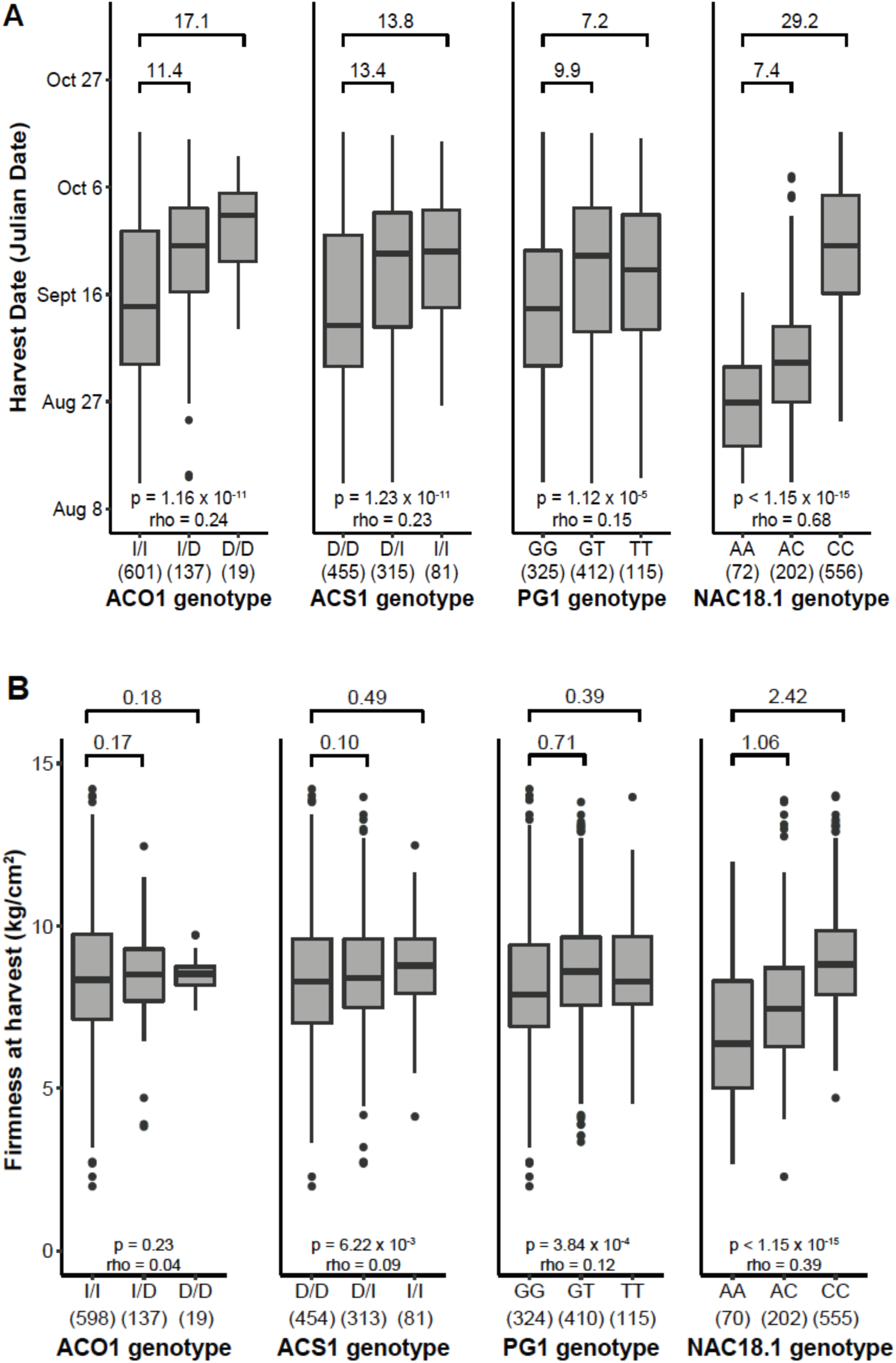
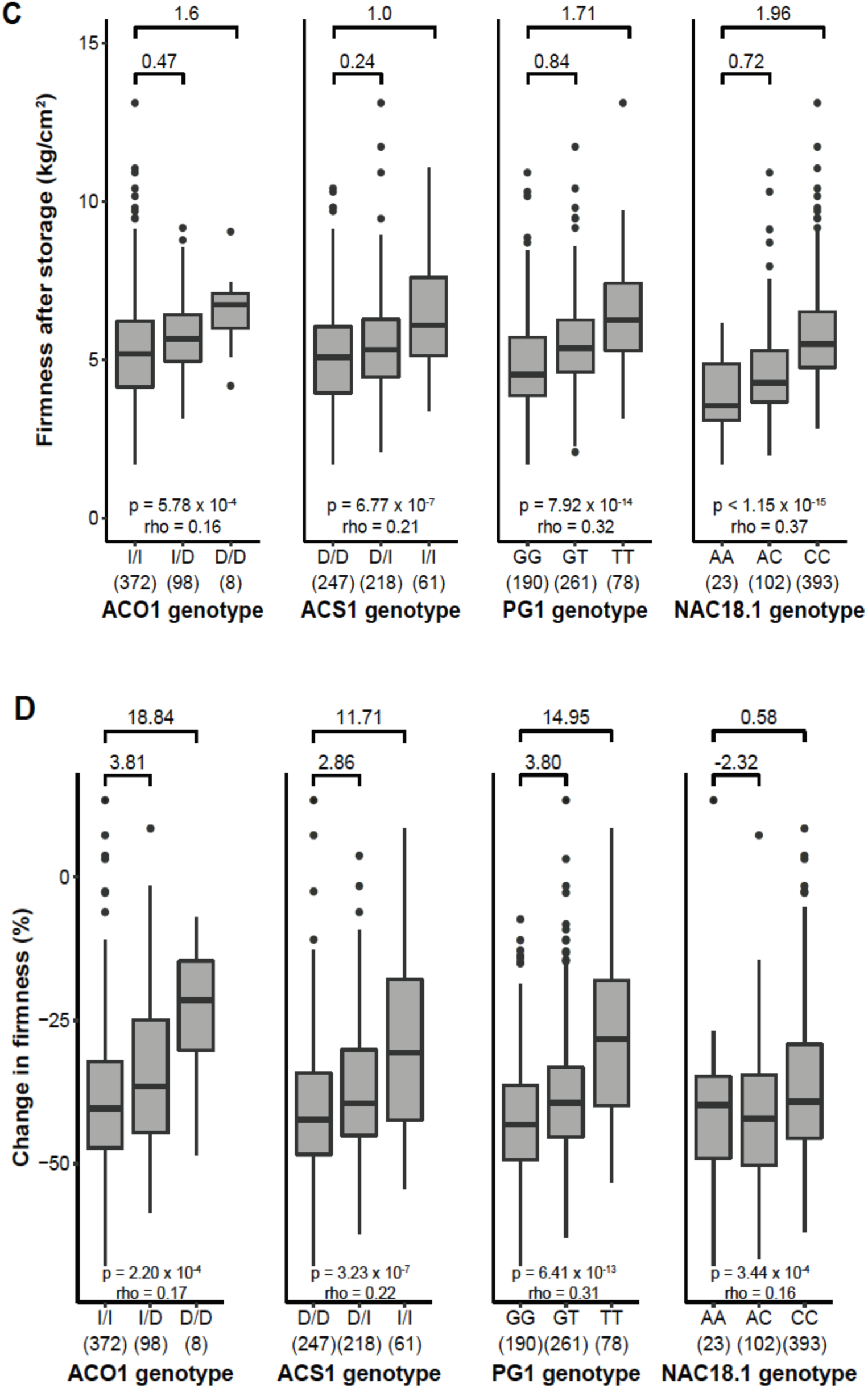
Genotype-phenotype correlations. Box plots display the variation in phenotypes across genotypic classes for four different firmness-related genetic markers. The phenotypes include (A) harvest date, (B) firmness at harvest, (C) firmness after storage, and (D) change in firmness (softening). Sample sizes are shown in parentheses under each genotypic class. The desirable genotype is on the right, while the undesirable genotype is on the left in every case. The P value and rho value are shown from performing a Spearman rank correlation test. The difference in phenotype values between genotypic classes are shown across the top of each plot.

**Figure S3:**
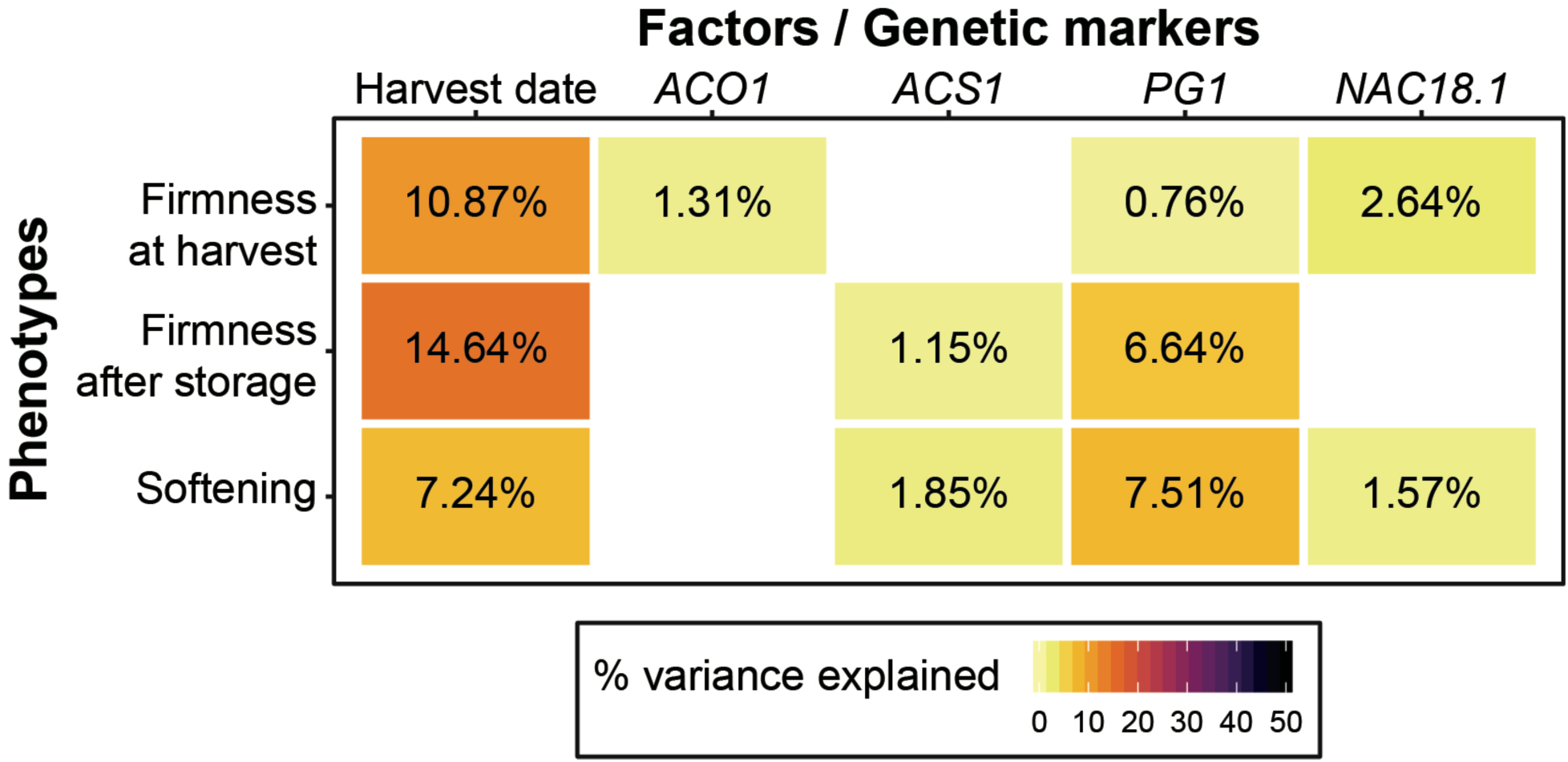
Prediction of firmness-related apple phenotypes. Four genetic markers and harvest date were included as factors in a type 2 ANOVA with three different phenotypes as outcomes. The proportion of the variance explained is shown in cases where a statistically significant result (P < 0.05) was observed.

**Figure S4:**
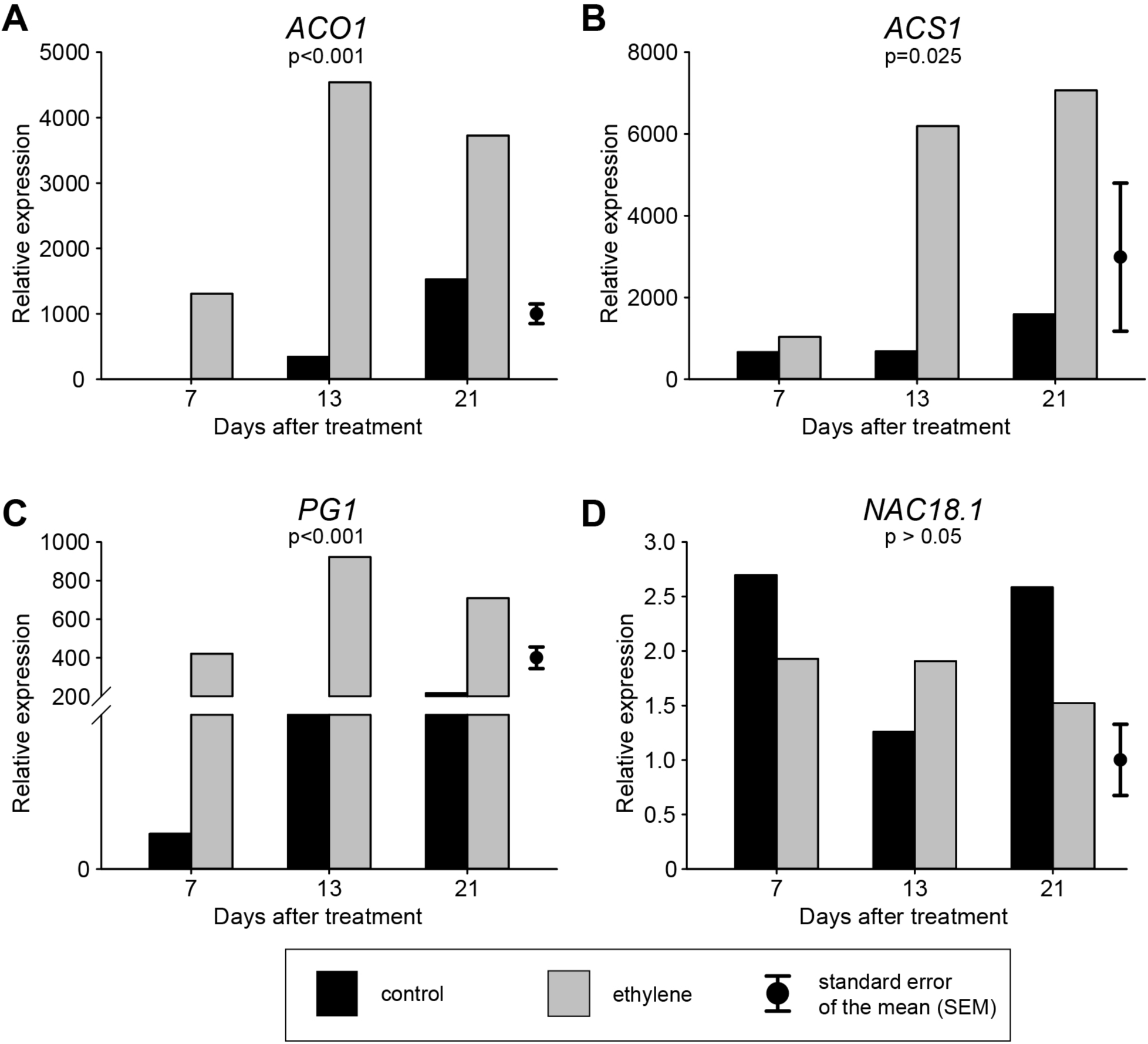
Relative expression of four firmness-related genes with and without exposure to ethylene over 3 weeks of storage. The ANOVAs testing for the effects of ethylene were significant (p < 0.05) for ACO1 (a), ACS1 (b) and PG1 (c), but not for NAC18.1 (d). Due to the balanced design, one standard error of the means (SEM) bar is used to represent the SEM for each population.

**Figure S5:**
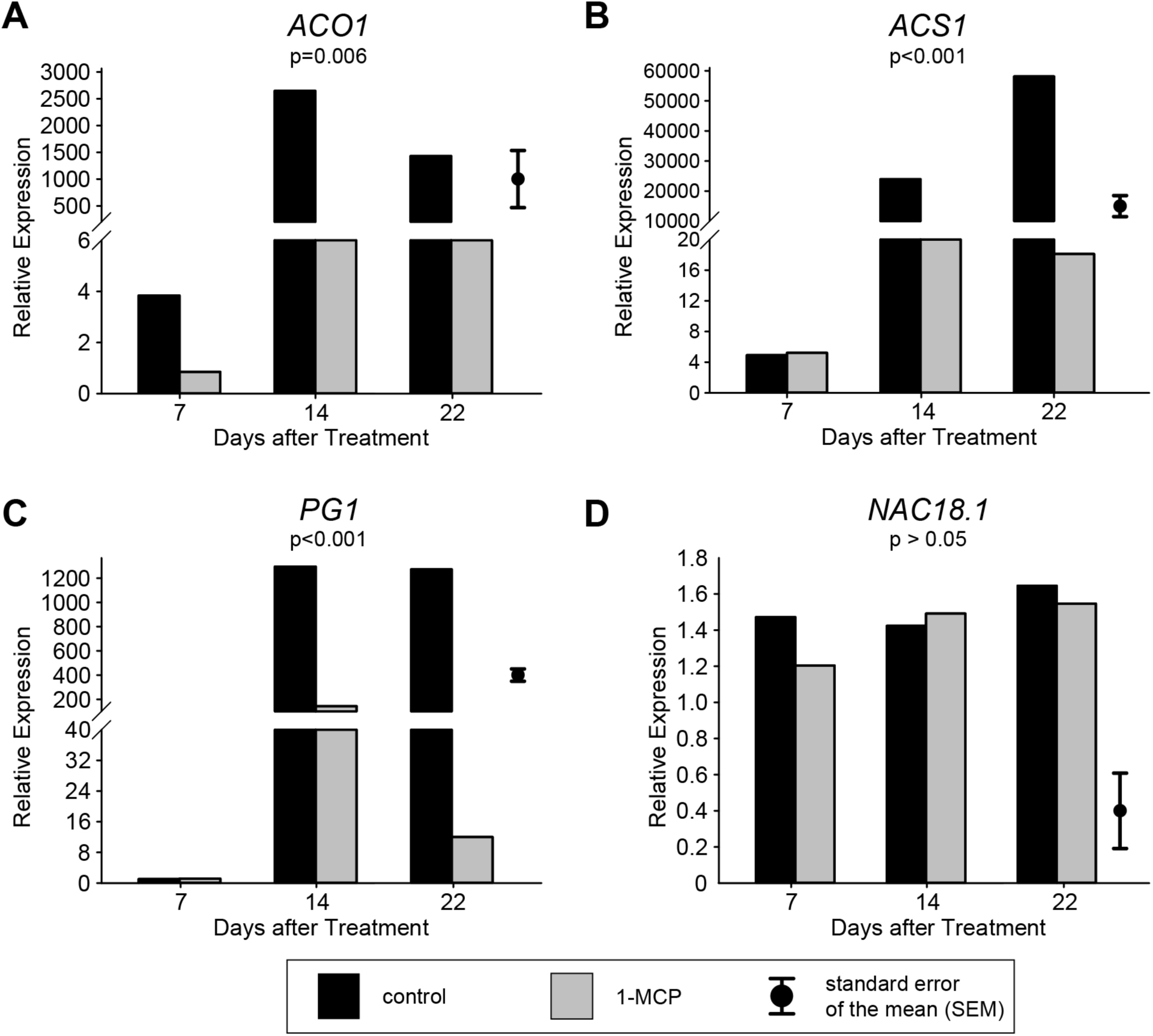
Relative expression of four firmness-related genes with and without exposure to the ethylene-inhibitor 1-MCP over 3 weeks of storage. The ANOVAs testing for the effects of 1- MCP were significant (p < 0.05) for ACO1 (a), ACS1 (b) and PG1 (c), but not for NAC18.1 (d). Due to the balanced design, one standard error of the means (SEM) bar is used to represent the SEM for each population.

**Figure S6:**
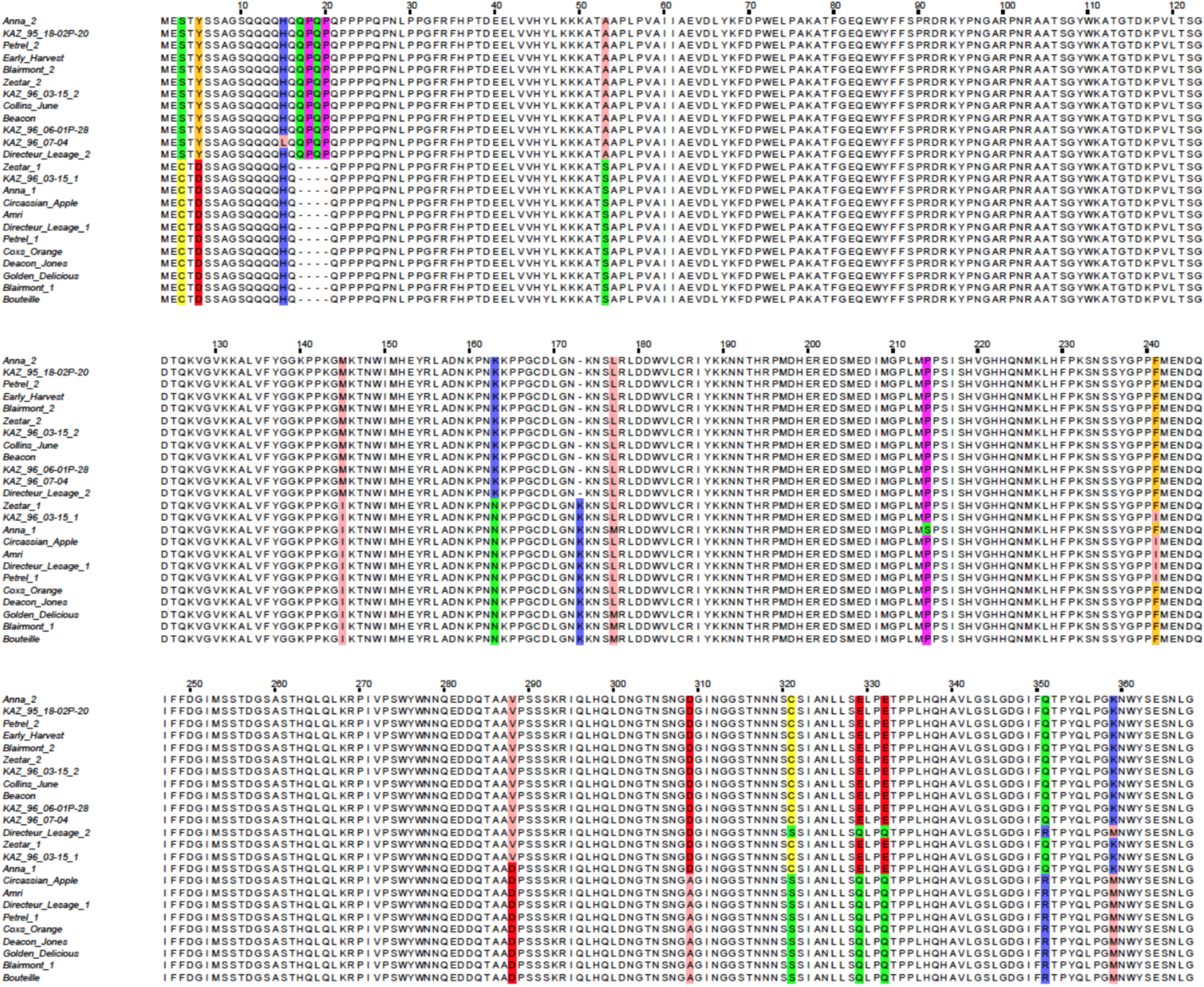
Amino acid sequence alignment of *NAC18.1*. Polymorphisms are highlighted in colour. Raw sequencing data is available in Table S4 in fasta format.

**Figure S7:**
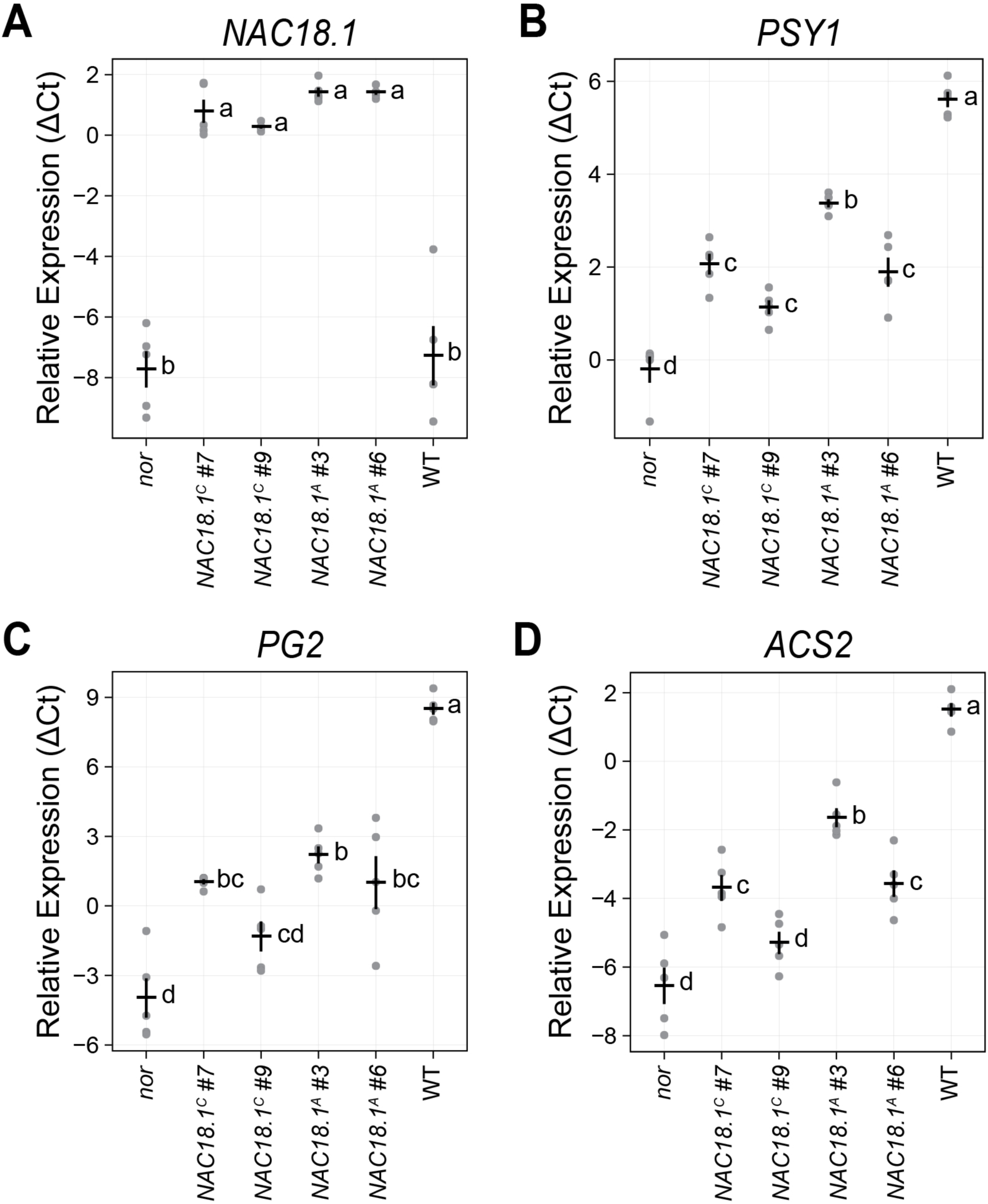
Ripening marker gene expression in transgenic tomatoes. Gene expression was evaluated as the difference in threshold cycle (ΔC_t_, log_2_ scale) by qRT-PCR using *RPL2* as a reference gene. The mean ± SE is superimposed in black over the raw values in gray (*n* = 5). Genotypes not sharing a letter (a-d) are statistically distinct by one-way ANOVA and Tukey’s HSD test (*p* < 0.05). (A) The *NAC18.1* transgene. (B) *PSY1*, encoding the phytoene synthase carotenoid biosynthetic gene. (C) *PG2*, encoding the major ripening-associated polygalacturonase involved in pectin hydrolysis. (D) *ACS2*, encoding 1-aminocyclopropane-1- carboxylate synthase, a ripening-associated isoform of the ethylene biosynthesis enzyme.

## Supplemental Table Legends

Table S1: Primer names and sequences.

Table S2: A list of sample IDs, sample names, genotypes and phenotypes.

Table S3: Efficiency values for gene expression assays of apple genes ACO1, ACS1, PG1 and NAC18.1.

Table S4: DNA sequences from 24 *NAC18.1* haplotypes in fasta format. Amino acid sequence alignment is visualized in Figure S6.

