## Supplementary figures and images for "Apple ripening is controlled by a NAC transcription factor"

### FigureS1.pdf

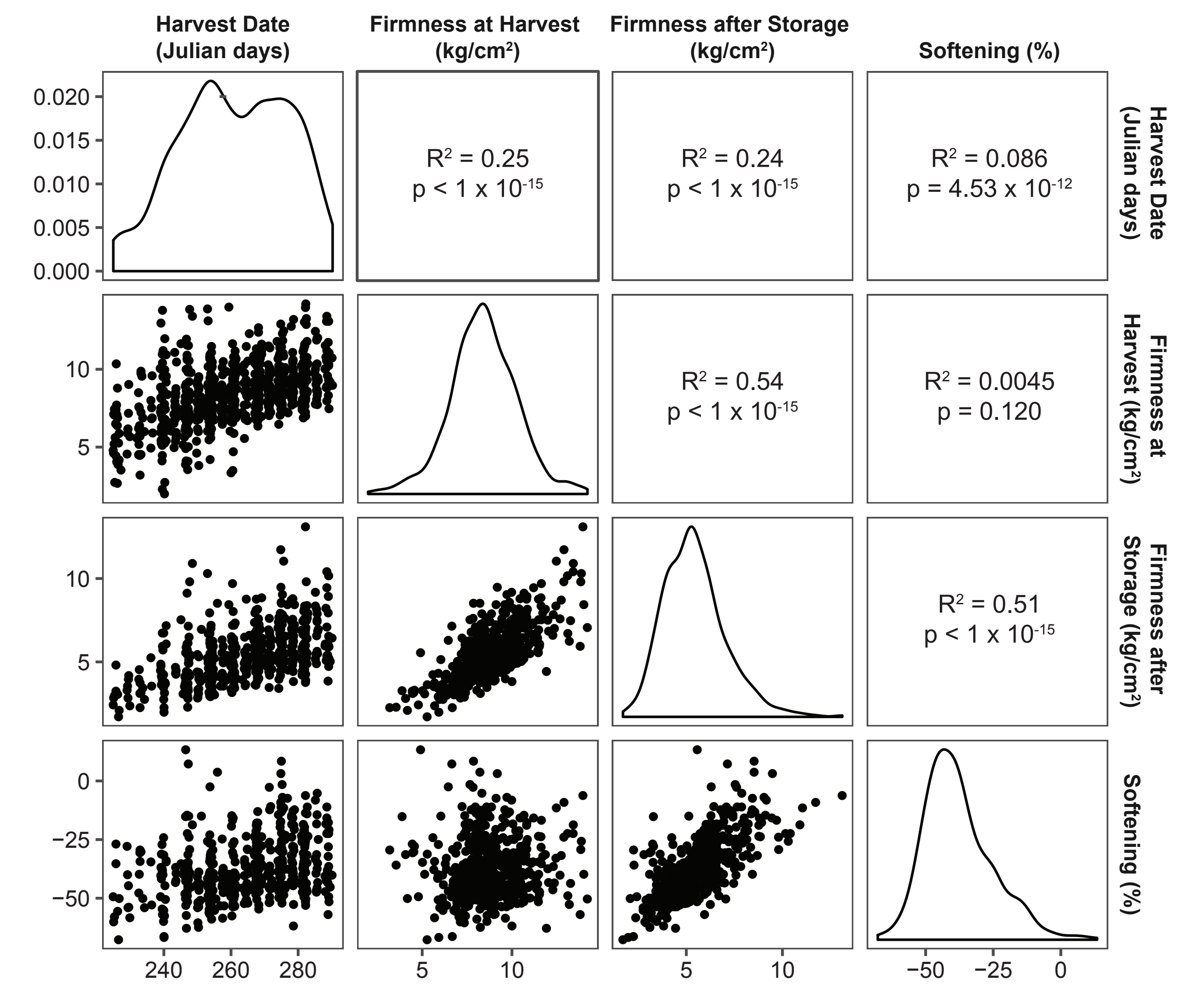

### FigureS2.pdf

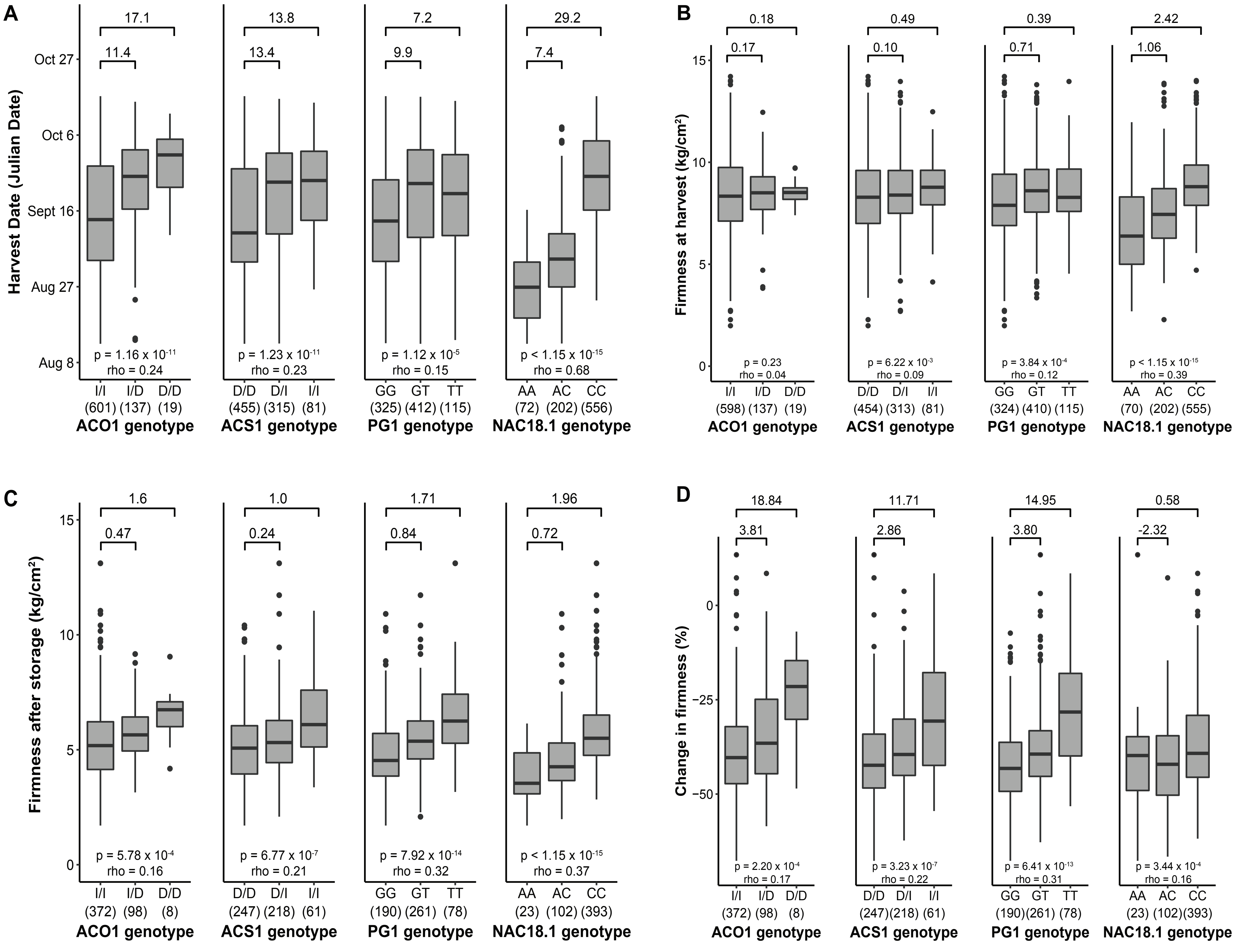

### FigureS3.pdf

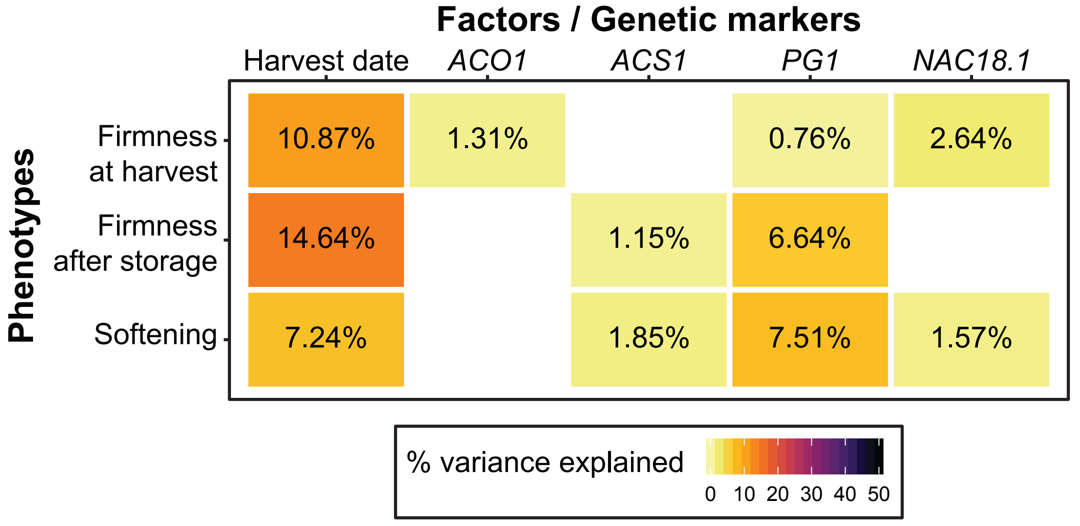

### FigureS4.pdf

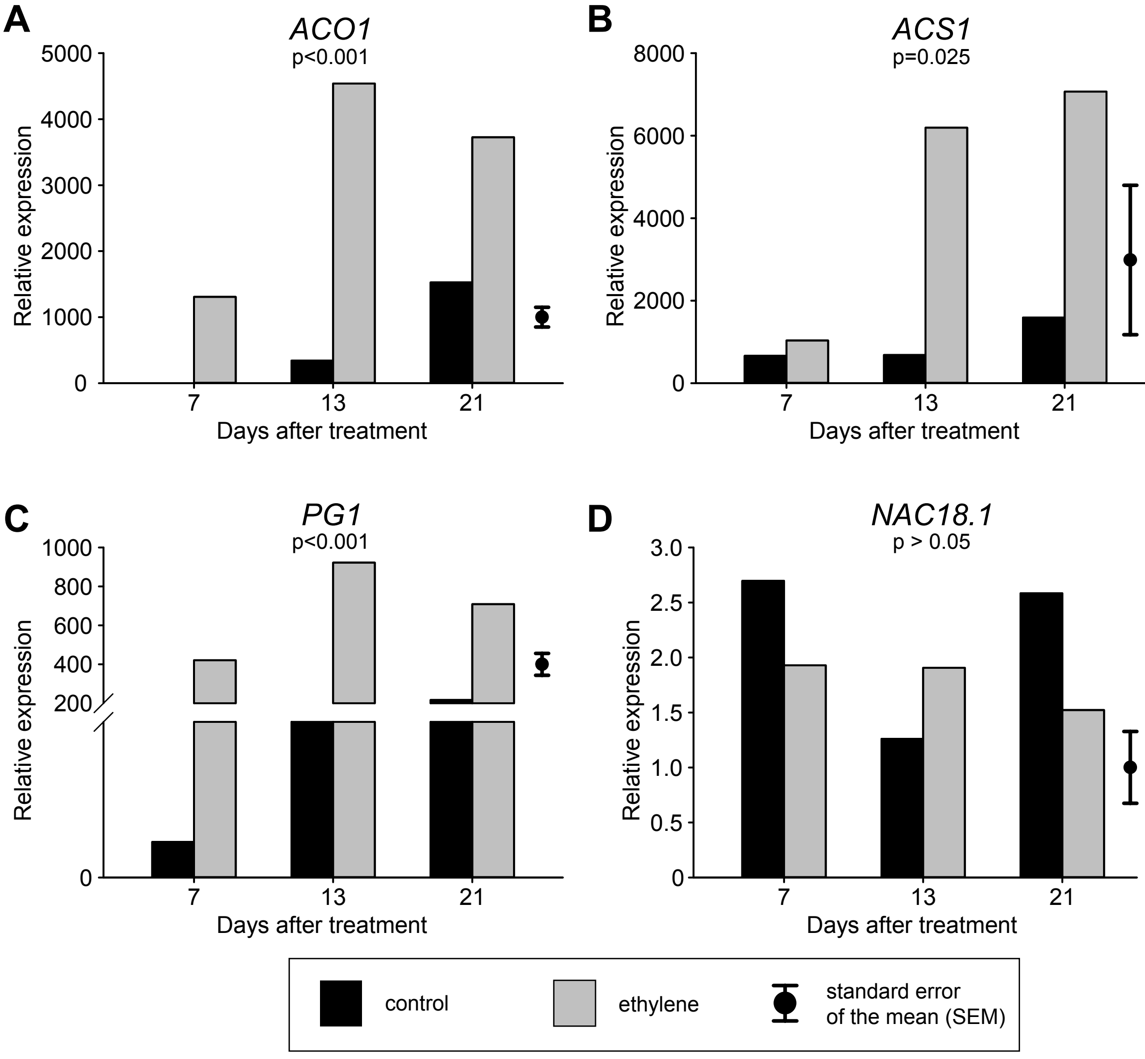

### FigureS5.pdf

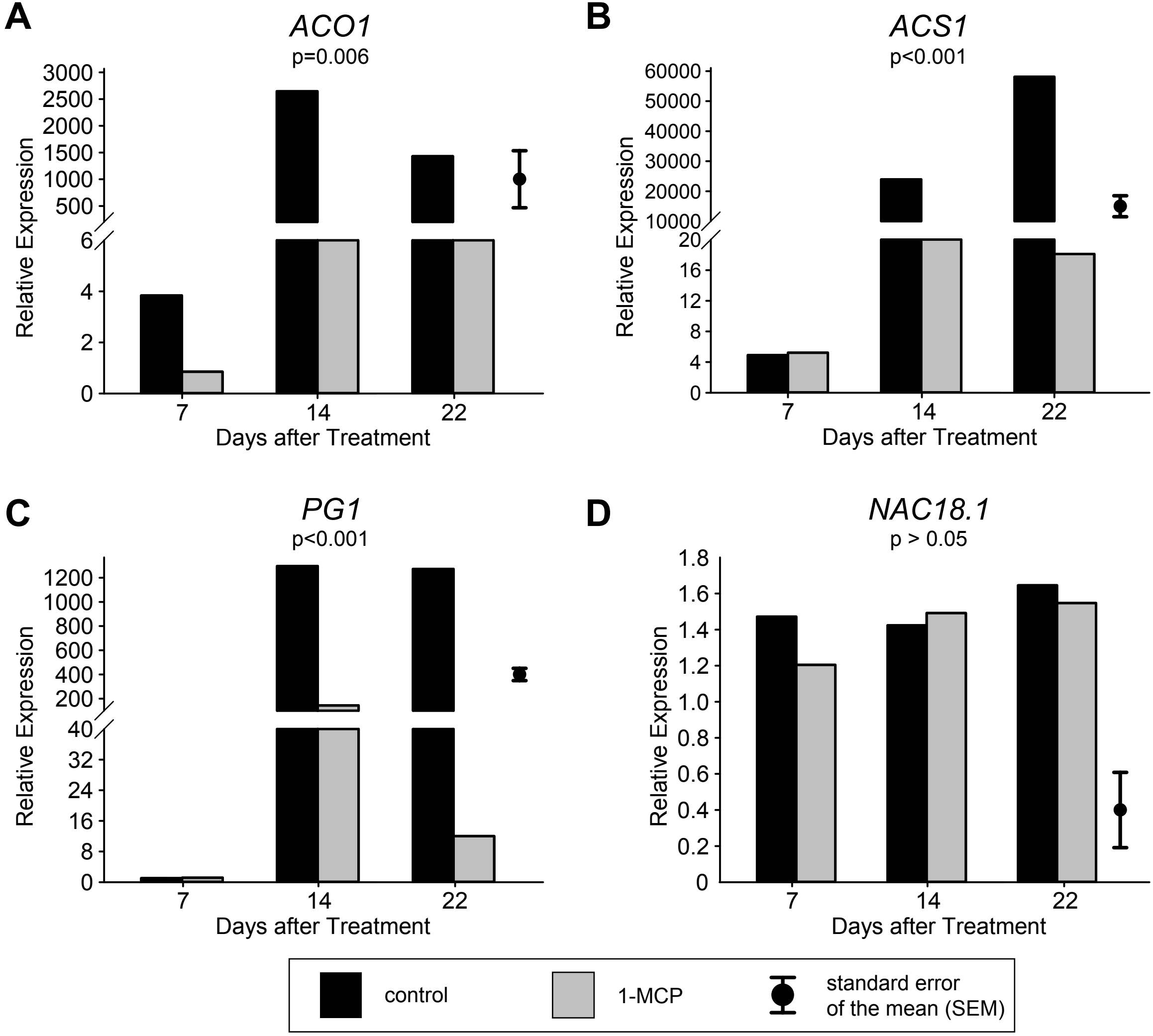

### FigureS6.pdf

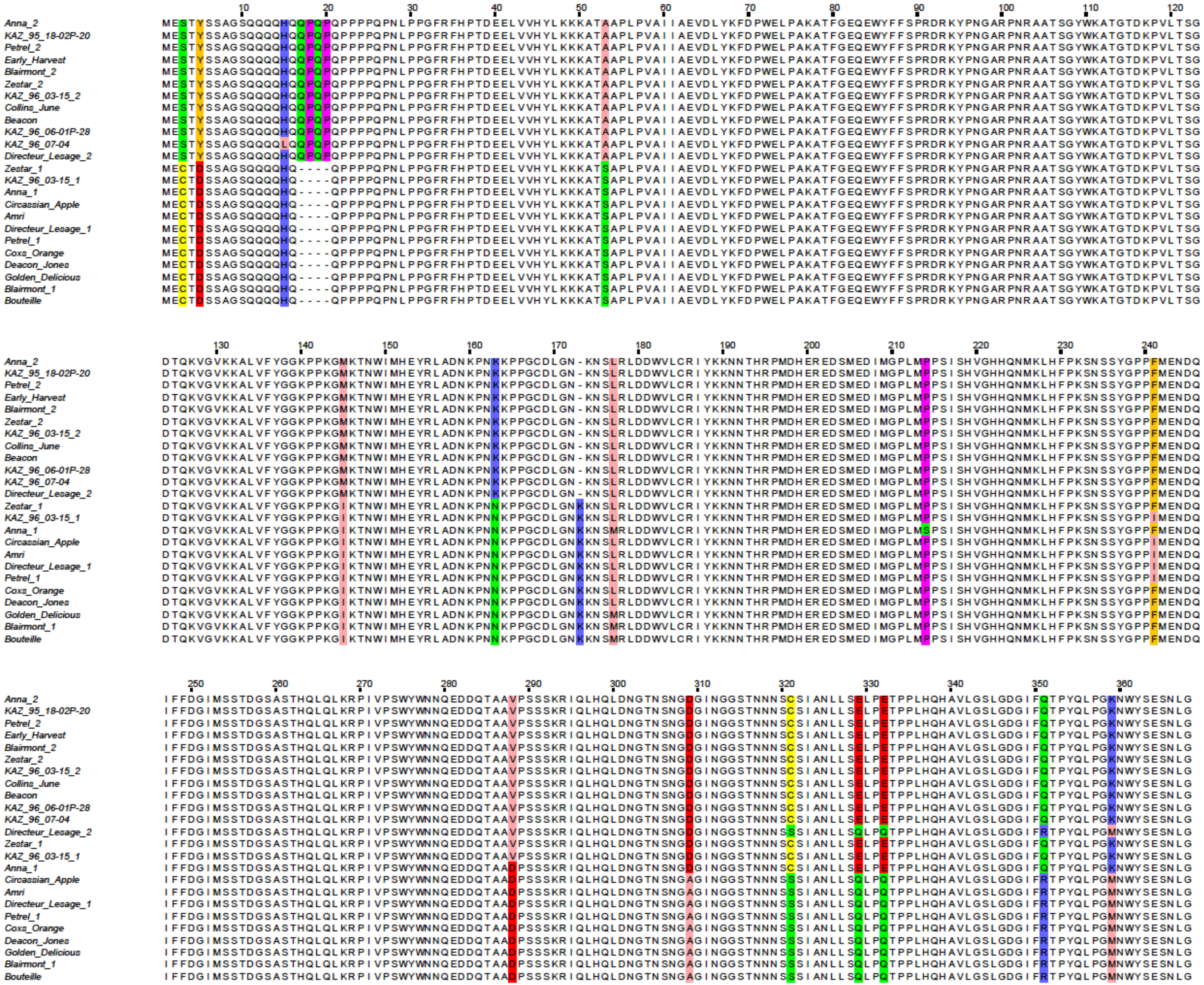

### FigureS7.pdf

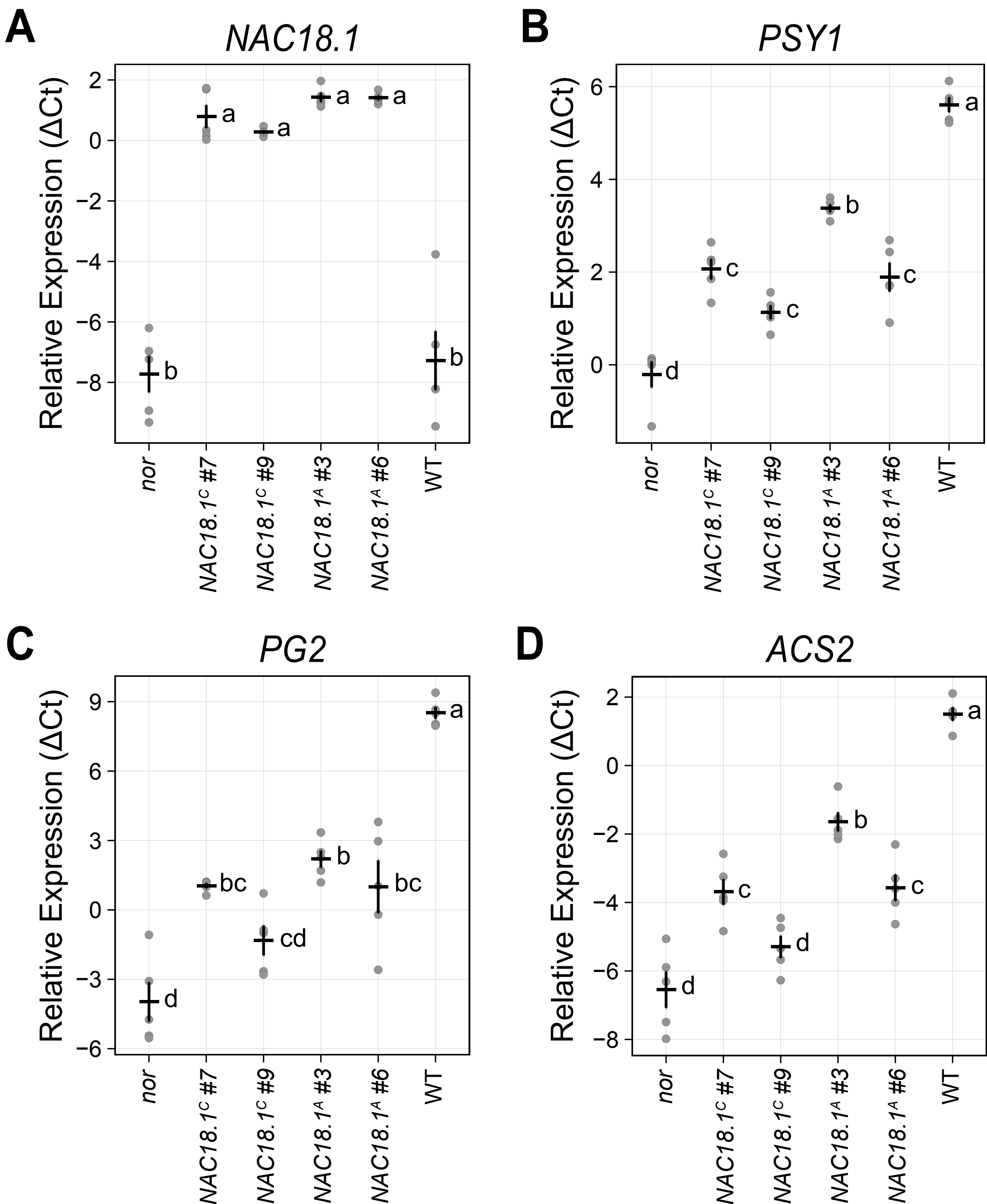
